# BET inhibition induces GDH1-dependent glutamine metabolic remodeling and vulnerability in liver cancer

**DOI:** 10.1101/2024.03.20.585859

**Authors:** Wen Mi, Jianwei You, Liucheng Li, Lingzhi Zhu, Xinyi Xia, Li Yang, Fei Li, Yi Xu, Junfeng Bi, Pingyu Liu, Li Chen, Fuming Li

## Abstract

Bromodomain and extra-terminal domain (BET) proteins, which function partly through MYC, are critical epigenetic readers and emerging therapeutic targets in cancer. Whether and how BET inhibition simultaneously induces metabolic remodeling remains unclear. Here we find that even transient BET inhibition by JQ-1 and other pan-BET inhibitors blunts liver cancer cell proliferation and tumor growth. BET inhibition decreases glycolytic gene expression but enhances mitochondrial glucose and glutamine oxidative metabolism revealed by metabolomics and isotope labeling analysis. Specifically, BET inhibition downregulates miR-30a to upregulate glutamate dehydrogenase 1 (GDH1) independent of MYC, which produces α-ketoglutarate for mitochondrial oxidative phosphorylation (OXPHOS). Targeting GDH1 or OXPHOS is synthetic lethal to BET inhibiton, and combined BET and OXPHOS inhibition therapeutically prevents liver tumor growth *in vitro* and *in vivo*. Together, we uncover an important epigenetic-metabolic crosstalk whereby BET inhibition induces MYC-independent and GDH1-dependendent glutamine metabolic remodeling that can be exploited for innovative combination therapy of liver cancer.

## Introduction

Primary hepatocellular carcinoma (HCC), the major type of liver cancer, is a leading cause of worldwide cancer-related death^1–3^. End stage liver cancer patients have limited treatment options due to the lack of druggable targets, and the treatment outcome is usually complicated by the heterogeneous tumor microenvironment^4^. Recent clinical trial show that only a small portion of patients initially respond to either molecular targeted therapy or combined anti-angiogenic therapy and immune checkpoint blockade, but eventually develop drug resistance^5, 6^. Therefore, it’s urgent to understand the drug resistance mechanisms and develop improved intervention strategies for liver cancer treatment.

One hallmark of cancer is deregulated metabolism, which contributes to tumor progression, metastasis and relapse through cancer cell intrinsic metabolic remodeling and metabolic interactions within tumor microenvironment^7–10^. HCC develops from transformed hepatocytes with features of metabolic remodeling^11^. For example, aberrant WNT signaling upregulates glutamine synthetase GLUL to increase glutamine levels and activate mTOR pathway to facilitate tumor progression^12^, loss of p53 activates SREBP-2-driven mevalonate pathway to promote liver tumorigenesis^13^. Moreover, loss of urea cycle enzymes in HCC renders the cancer cells auxotrophic for arginine^14^, and ASS1 downregulation enables cancer cells to accumulate aspartate for pyrimidine synthesis necessary for cell proliferation^15^. Recently, we demonstrated that gluconeogenic gene Fructose-1,6-Bisphosphatase (FBP1) is universally silenced during liver tumorigenesis, and hepatic FBP1 loss in mice disrupts liver metabolism and promotes liver tumor progression, through a hepatic stellate cell (HSC) senescence secretome and extracellular vesicle-mediated communication between hepatocytes and natural killer cells^16, 17^. While these studies have identified different metabolic enzymes and pathways modulating liver tumor initiation and progression, how liver cancer cells undergo metabolic adaptations in response to specific drug treatment has not been fully explored.

Bromodomain and extra terminal domain (BET) proteins, which comprises BRD2, BRD3, and BRD4 and the testis-restricted BRDT, are epigenetic readers containing two tandem bromodomains (BD1 and BD2), an extra terminal domain (ET), and a C-terminal domain (CTD)^18^. By recognizing acetylated lysine of histone and non-histone proteins, BET proteins act as scaffolds to recruit many other proteins to promoters and enhancers (especially at the super enhancers) of target genes and to facilitate the transcription. Notable BET target genes include *MYC, BCL2, BCL6, CDK4* and *CDK6*, which are involved in cell cycle progression, cell survival and other biological processes. Among these, the oncogene MYC is also a master regulator of cellular metabolism. Overexpressed MYC, observed in several types of cancer, activates target genes of glycolysis, mitochondrial biogenesis/ oxidative phosphorylation (OXPHOS), glutamine metabolism, *de novo* nucleotide synthesis and many other metabolic pathways^19, 20^. Given the overexpression of BET proteins in cancer, previous studies have shown the promise of targeting BET proteins as an intervention strategy in hematologic malignancies, breast cancer^21^, prostate cancer^22^, etc. Consistently, one study showed that the pan-BET inhibitor JQ-1 reduces liver fibrosis^23^, or liver tumor burden in a mouse model of non-alcoholic steatohepatitis (NASH)-HCC^24^. In the meanwhile, different classes of pan-BET inhibitors (BETi) are now in clinical trials, which confirm their antitumor potential^25, 26^. However, the efficacy of BET inhibition as monotherapy is limited, making it necessary to develop combination therapies with other anticancer agents, currently including kinase inhibitors, epigenetic drugs, immune modulators and hormone therapy^26^. Since cancer cells dynamically undergo metabolic remodeling during drug treatment, how BET inhibition induces MYC-dependent or independent metabolic adaptations in cancer cells remains unclear. Addressing this question will add more mechanistic insights into how epigenetic-metabolic crosstalk contributes to tumor growth, and more importantly uncover metabolic vulnerabilities that could be exploited to enhance therapeutic efficacy of BET inhibition in cancer.

In this study, we show that BET inhibition induces MYC-independent and GDH1-dependent glutamine metabolic remodeling in liver cancer, and that targeting the GDH1-OXPHOS metabolic axis is “synthetic lethal” to BET inhibition, which limits liver tumor progression in different mouse models. Our study thus provides an innovative combination therapy for liver cancer treatment.

## Results

### Transient BET inhibition blunts HCC cell growth and induces differentiation

Among the BET family, BRD4 is the best-studied member to regulate tumorigenesis through cancer cell autonomous and non-autonomous mechanisms^18, 26^. We set out to determine the correlation of individual BET gene expression with liver cancer patient survival. Analysis of TCGA dataset demonstrated that elevated BRD2, BRD3 and BRD4 transcript levels negatively correlated with patient survival (Fig. 1a). Meanwhile, shRNA-mediated knockdown of BRD2 and BRD4 in the liver cancer cell line Huh7 significantly slowed down cell proliferation, suggesting both BRD2 and BRD4 are important for liver cancer cell growth (Fig. 1b, c). To pharmacologically target BET proteins, we applied the well-established pan-BET inhibitor JQ-1^27^ to treat Huh7 and Hep3B cells, two most sensitive cell lines according to the “The Genomics of Drug Sensitivity in Cancer Project” (https://www.cancerrxgene.org) (Extended Data Fig. 1a). JQ-1 dose-dependently decreased growth without inducing dramatic death in both cell lines. Similarly, another orally bioavailable and potent pan-BET inhibitor ABBV075^28^ also blunted Huh7 and Hep3B cell growth in a dose-dependent manner (Extended Data Fig. 1b-d). Notably, even after transient (48 hour) JQ-1 exposure, both Huh7 and Hep3B cells exhibited much slower proliferation in cell growth assay (Extended Data Fig.1e), and formed fewer colonies in clonogenicity assay compared to vehicle-treated cells (Fig. 1d, e). In a serum-free suspension culture condition to enrich CSC-derived spheres^29^, transient JQ-1 pre-treated Huh7 and Hep3B cells gave rise to much smaller and fewer spheres, further supporting reduced tumorigenicity potential (Fig.1f, g). Accordingly, when control and BETi-exposed Huh7 cells were subcutaneously transplanted to immune compromised mice, the resultant tumor growth was profoundly dampened, as supported by decreased tumor volume change (Fig. 1h).

**Fig. 1.**
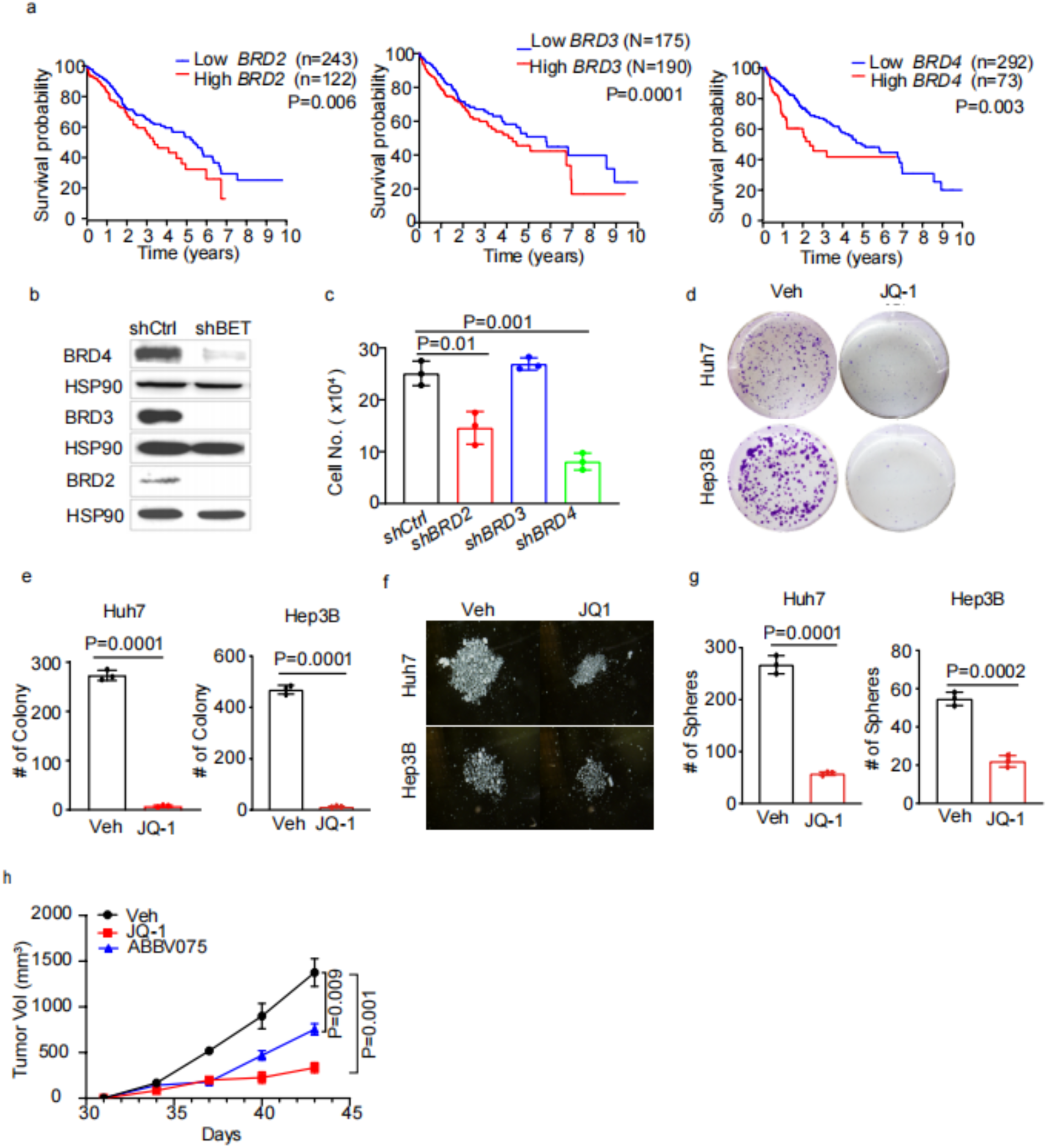
BET inhibition blunts HCC cell growth and induces differentiation. **a**, Kaplan–Meier overall survival plots stratified by *BRD2, BRD3* and *BRD4* mRNA levels from TCGA HCC database. A log-rank Mantel–Cox test was performed between the groups. **b**, Western blot analysis of indicated proteins from control and BET-knock down Huh7 cells. **c** Cell number quantification after growing control and BET-knock down Huh7 cells for 72 hours **d**, Representative clonogenicity assays from replated Huh7 and Hep3B cells after vehicle and JQ-1 (0.5 μM) pretreatment for 48 hours. **e**, Colony number quantification of replated Huh7 and Hep3B cells after vehicle and JQ-1 (0.5 μM) pretreatment for 48 hours. **f, g**, Representative Huh7 and Hep3B sphere cultures (h) and quantification (i) from vehicle and JQ-1 pretreatment groups. **h**, Huh7 xenograft tumor growth curve from vehicle (n=4) and JQ-1 (n=6) or ABBV (n=6) pre-treatment groups. All statistical graphs except (**a**) show the mean ± s.e.m. Except (**a**), *P* values were calculated using a two-tailed Student’s t-test. All experiments were performed in biological triplicate.

BETs contain two bromodomains (BDs), and it’s been shown that BD1-specific inhibitors phenocopy the effects of pan-BET inhibitors whereas BD2 inhibitors are predominantly effective in inflammatory and autoimmune disease models^33^. We treated Huh7 cells with BD1 (GSK778) or BD2 (GDK046) inhibitors, and found that Huh7 cells were only sensitive to BD1 inhibitor but refractory to BD2 inhibitor (Extended Data Fig.1f), suggesting that BD1 mediates the oncogenic functions of BETs in liver cancer. Collectively, these results indicate that even transient BET inhibition is sufficient to blunt liver cancer cell proliferation and tumor growth.

### BET inhibition alters glucose and glutamine metabolic gene expression

To address whether BET inhibition results in metabolic remodeling in liver cancer cells, RNA-sequencing was performed to compare the transcriptome and metabolic gene expression in control and transient JQ-1-treated Huh7 cells. This profiling uncovered significantly altered mRNAs (padj< 0.05, more than 2 fold change), including 1743 genes upregulated and 2027 genes downregulated loci. Gene set enrichment analysis (GSEA) further uncovered top-ranked gene sets of glycolysis, oxidative phosphorylation (OXPHOS), purine metabolism and pyrimidine metabolism that were significantly altered by JQ-1 exposure (Fig.2a and Extended Data Fig.2a). Consistently, KEGG analysis of metabolic pathways identified glycolysis as one of top-ranked pathways with distinct gene expression pattern (Fig.2b). Indeed, a volcano plot of metabolic gene expressions clearly showed that many glycolytic gene transcripts were downregulated (Fig.2c and Extended Data Fig.2b), and both mRNA and protein levels of representative glycolytic genes (*GLUT1*, *HK2*, *LDHA*) were validated by quantitative reverse-PCR (Q-PCR) and western blot analysis, respectively (Fig. 2d and Extended Data Fig. 2c).

**Fig. 2.**
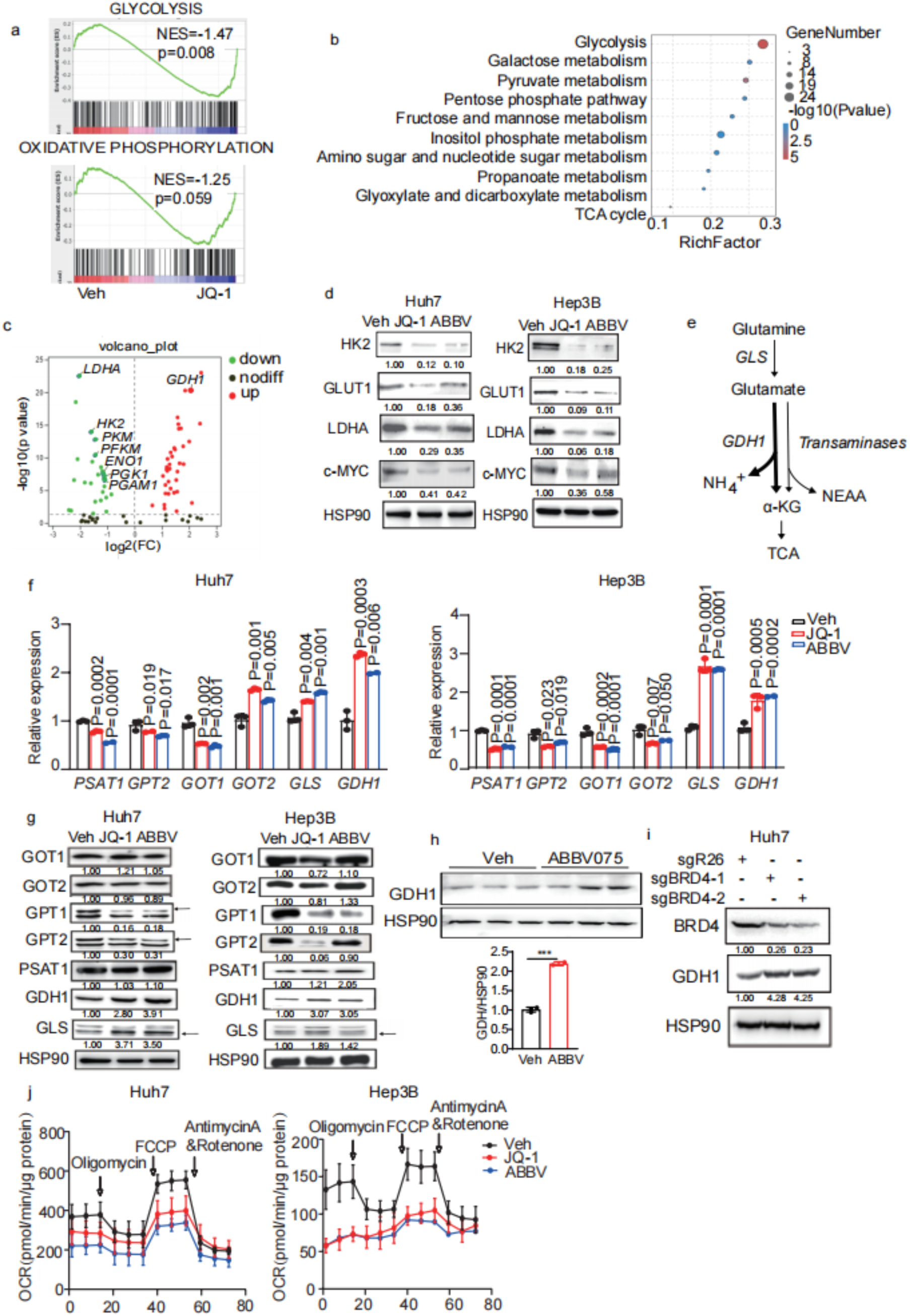
BET inhibition alters glucose and glutamine metabolic gene expression. **a**, GSEA for glycolysis and oxidative phosphorylation from vehicle and JQ-1 groups based on RNA-sequencing data. **b**, Top-ranked KEGG metabolic pathways from vehicle and JQ-1 groups based on RNA-sequencing data. **c**, A volcano plot of indicated metabolic gene expression based on RNA-sequencing data. **d**, Western blot analysis of glycolytic enzymes from vehicle and BETi-treated Huh7 and Hep3B cell lysates. HSP90 is used a loading control. **e**, A scheme of glutaminolysis. **f**, **g**, Q-PCR (**f**) and western blot (**g**) analysis of glutamine metabolic gene expression in BETi-treated Huh7 and Hep3B cells. **h**, Western blot analysis and quantification of GDH1 expression in Huh7 xenograft tumor model using ABBV075. n=3 tumors for each group. **i,** Western blot analysis of GDH1 from sgCtrl and sgBRD4 Huh7 cell lysates**. j**, Seahorse assays of vehicle and BETi-treated Huh7 and Hep3B cells. All statistical graphs show the mean ± s.e.m. *P* values were calculated using a two-tailed Student’s t-test. All experiments were performed in biological triplicate.

Glutaminolysis provides the cells with nitrogen source, non-essential amino acids (NEAA) including alanine, aspartate and proline, as well as α-ketoglutarate (α-KG) which can replenish the tricarboxylic acid (TCA) cycle^34^. Specifically, mitochondrial glutamine is converted to glutamate by glutaminase (GLS1), and glutamate is then catabolized either through GDH1 to produce α-KG, or through different transaminases to generate NEAA and α-KG (Fig.2e). Since RNA-sequencing revealed *GDH1* upregulation by JQ-1 treatment in Huh7 cells, we then experimentally compared glutaminolysis-related gene expressions. Both mRNA and protein levels of GLS1 and GDH1 were consistently elevated by JQ-1 or ABBV075 treatment in Huh7 and Hep3B cells, while transaminase-coding genes exhibited heterogenous changes in mRNA or protein levels. For example, GPT1 and GPT2 expression were decreased, while PSAT1, GOT1 and GOT2 levels remained largely unchanged (Fig.2f, g). Similarly, GDH1 upregulation was also detected in ABBV075-treated Huh7 xenografts, as well as Huh7 cells with CRISPR-mediated BRD4 depletion (Fig.2i).

In addition to glycolysis and glutaminolysis genes, we also asked whether BET inhibition may directly impact mitochondrial gene expression. Protein levels of TFAM, a critical transcriptional regulator of mitochondrial biogenesis, was reduced by BETi treatment (Extended Data Fig.2d). Consistently, TFAM targets encoding mitochondrial electron transport chain (ETC) subunits (*MT-ND1*, *NDUFA11*, *ATP5D*, *ATP8*, *UCP2* and *COX17*) exhibited reduced transcription by BETi (Extended Data Fig.2e). Importantly, BETi-treated Huh7 and Hep3B cells had reduced oxygen consumption rate (OCR) in seahorse assays, accompanied with decreased basal and maximal respiration, as well as ATP production, indicating defective mitochondrial function (Fig.2j and Extended Data Fig.2f). Together, we concluded that BET inhibition potentially elicits metabolic remodeling in liver cancer cells.

### BET inhibition enhances glutamine derived TCA anaplerosis and shift energy supply to TCA cycles

To profile the metabolic remodeling process upon BET inhibition, we performed metabolomics analysis on vehicle and JQ-1 treated Huh7 cells. This profiling uncovered top-ranked and enriched metabolic pathways, including glycolysis, glutamine/glutamate metabolism, glutathione metabolism, taurine and hypotaurine metabolism and others (Extended Data Fig. 3a, b). In line with the elevated GLS1 and GDH1 expression, abundance of glutaminolysis and TCA cycle intermediate metabolites (glutamate, α-KG, aspartate, succinyl CoA and isocitrate) was increased upon JQ-1 treatment, indicating enhanced glutamine anaplerosis for TCA cycle (Fig.3a). Notably phosphorylation of the energy sensor AMPK (pAMPK) remained unchanged by JQ-1 treatment, collectively suggesting that BETi-exposed cells are able to maintain energy homeostasis despite impaired growth (Extended Data Fig. 3c).

To further validate above results and determine the effects of BETi on glucose and glutamine metabolic remodeling, we sought to measure relevant metabolic fluxes using isotope tracing techniques in vehicle and JQ-1-treated Huh7 cells. Specifically, we cultured cells with uniformly labeled glucose ([U-^13^C] glucose) and measured ^13^C enrichment of intracellular metabolites by mass spectrometry. [U-^13^C]glucose derived M+3 pyruvate (containing 3 ^13^C carbons) undergoes oxidative decarboxylation to generate [1,2-^13^C] acetyl-CoA (CoA, coenzyme A) and subsequently M+2 labeled citrate. We observed the pyruvate decarboxylation flux increased (Fig. 3c), likely to compensate the energy supply loss due to the reduced glycolysis activity. Meanwhile, pyruvate could also enter TCA via pyruvate carboxylase (PC)-mediated carboxylation flux to supply TCA substrate (Fig. 3b). The flux modestly decreased as demonstrated by decreased M+3 malate (Fig. 3c).

**Fig. 3.**
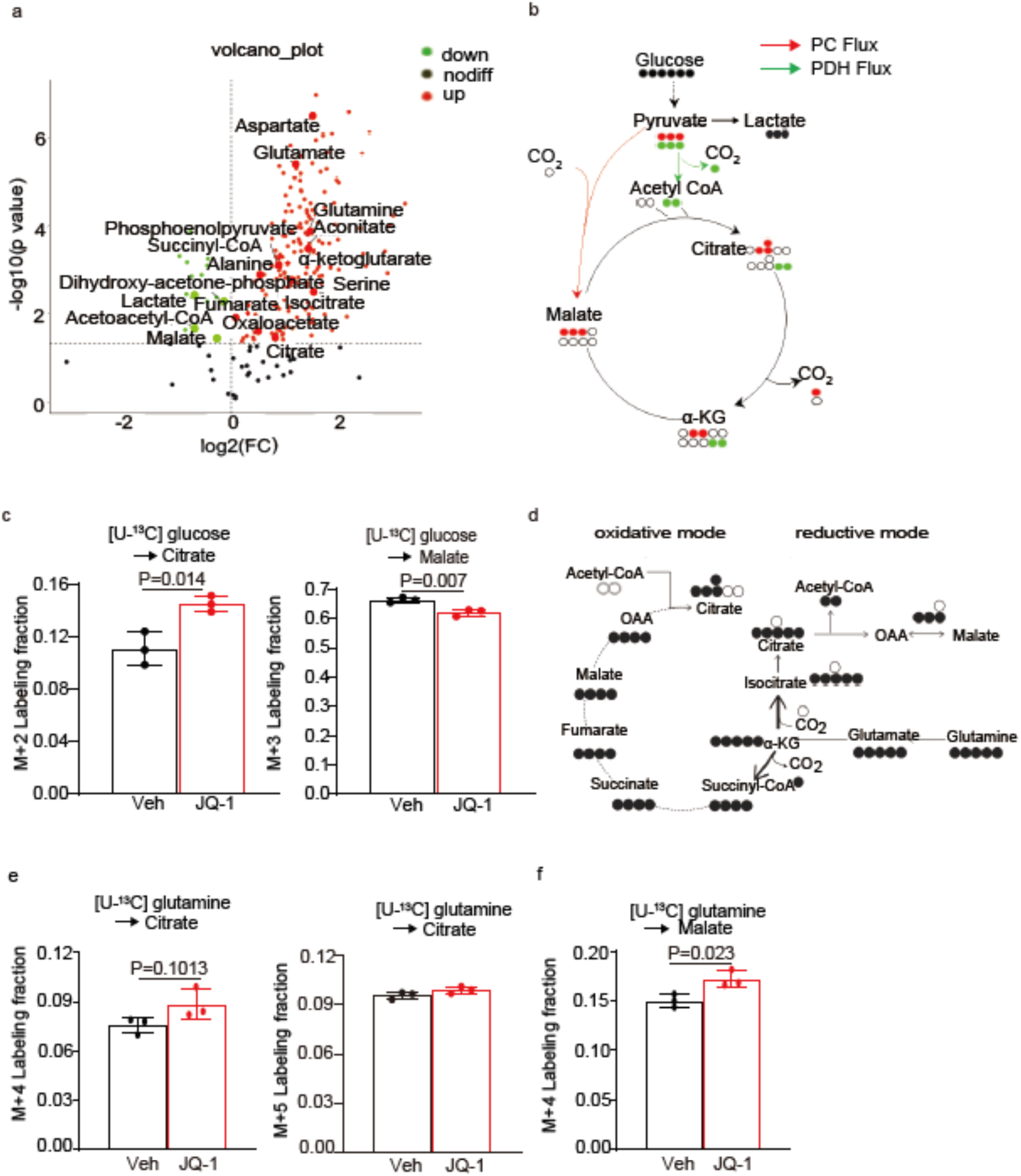
BET inhibition induces glutamine metabolic remodeling. **a**, A volcano plot of indicated metabolite abundance from targeted metabolomics of vehicle and JQ-1 treated Huh7 cells. **b**, A scheme of glucose labeling followed by PC or PDH-dependent metabolism to synthesize citrate and malate. **c**, Mass isotopologue analysis of M+2 citrate and M+3 malate in vehicle and JQ-1 pre-treated Huh7 cells cultured with U-^13^C glucose. **d**, A scheme of glutamine labeling followed by oxidative or reductive metabolism to synthesize citrate. **e**, Mass isotopologue analysis of M+5 or M+4 citrate in vehicle and JQ-1 pre-treated Huh7 cells cultured with U-^13^C glutamine. **f**, Mass isotopologue analysis of M+4 malate in vehicle and JQ-1 pre-treated Huh7 cells cultured with U-^13^C glutamine. All statistical graphs show the mean ± s.e.m. *P* values were calculated using a two-tailed Student’s t-test. All experiments were performed in biological triplicate.

We also cultured cells with uniformly labeled glutamine ([U-^13^C]glutamine) to probe glutamine related metabolic fluxes. [U-^13^C] glutamine-derived α-KG enters TCA cycle and undergoes oxidative metabolism (oxidative mode indicating energy supply) where TCA cycle metabolites including malate and citrate are M+4 labeled. Alternatively, α-KG undergoes reductive carboxylation (reductive mode indicating reductive biosynthesis) where α-KG is converted to M+5 isocitrate and finally to M+5 citrate (Fig.3d). Notably, as a function of the increased α-KG to citrate ratio^35^, reductive glutamine metabolism is essential to support growth of tumor cells with mitochondrial defects or maintain redox homeostasis during anchorage-independent growth^36–39^. Flux analysis revealed that JQ-1 exposure resulted in increased labeling fraction of citrate M+4 and malate M+4 to different extents, while labeling fraction of citrate M+5 remained comparable(Fig. 3e,f). This labeling result indicates enhanced oxidative glutamine metabolism, while reductive glutamine metabolism was minimally affected, and is consistent with higher GLS and GDH1 transcript levels upon BET inhibition.

Together, both metabolomics profiling and isotope labeling results suggest that BET inhibition induces glutamine derived TCA anaplerosis and the energy supply shift to TCA cycles to support cell growth.

### GDH1-dependent glutamine metabolism maintains cancer cell viability after BET inhibition

We have now shown that BET inhibition upregulated GDH1 expression and simultaneously decreased certain mitochondrial transaminase (GPT2) expression, whereas abundance of α-KG, a common product of these enzymes, was not reduced but increased instead. We thus reasoned that GDH1 upregulation preserves α-KG levels for TCA cycle to maintain energy metabolism. If this is case, then GDH1 would be required for the cell growth and/or viability after BET inhibition. To test this, we knocked down GDH1 in Huh7 cells, and then treated control (*shCtrl*) and *GDH1* knockdown (*shGDH1*) cells with JQ-1 (Fig. 4a). As shown in Fig. 4b and c, compared to control cells, GDH1-deficient Huh7 cells were extremely sensitive to JQ-1 treatment and dramatic cell death was detected by PI/Annexin V staining and flow cytometry.

**Fig. 4.**
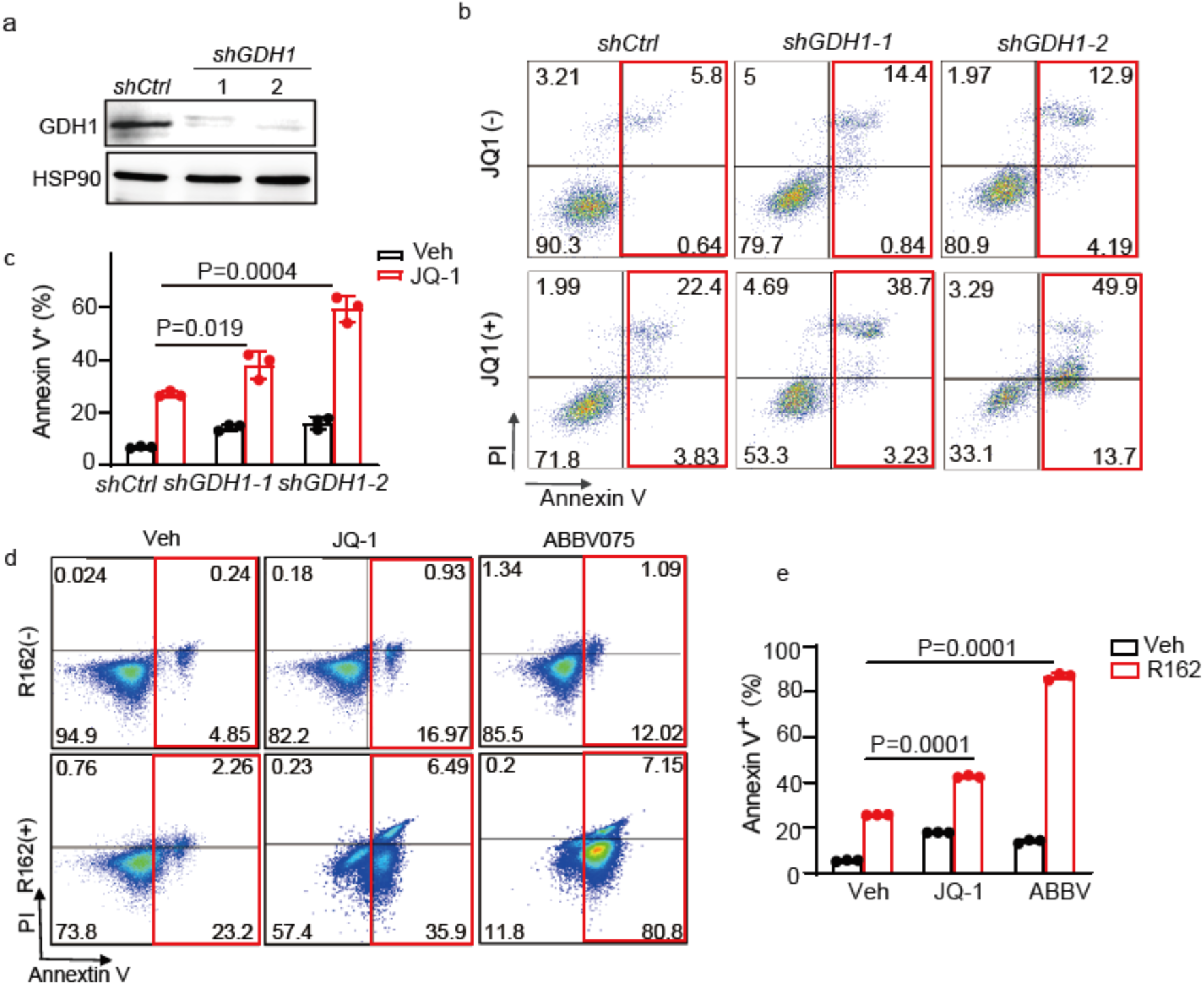
*GDH1*-dependent glutamine metabolism maintains cell viability after BET inhibition. **a**, Western blot analysis of *GDH1* expression in control (*shCtrl*) and *GDH1* knock down (*shGDH1*) Huh7 cells. HSP90 is used as a loading control. **b**, Representative flow cytometry plot of PI and annexin V staining of Huh7 cells from indicated groups. **c**, Statistic analysis of the percentage of annexin V^+^ cells of indicated groups in (**b**). **d**, Representative flow cytometry plot of PI and annexin V staining in the vehicle and JQ-1/R162 treated Huh7 cells. **e**, Statistic analysis of the percentage of annexin V^+^ cells of indicated groups in (**d**). All statistical graphs show the mean ± s.e.m. *P* values were calculated using a two-tailed Student’s t-test. All experiments were performed in biological triplicate.

It was previously shown that a small molecule inhibitor R162 can abrogate GDH1 enzyme activity and dramatically decrease intracellular α-KG levels in lung and breast cancer cells^40^. We then asked whether R162-treated Huh7 cells would similarly be vulnerable to BETi. Although R162 itself didn’t induce profound cell death in Huh7 cells, combined R162 and JQ-1 or ABBV075 exposure remarkably induced apoptosis (Fig. 4d, e). As expected, pharmacologically targeting GLS1 using CB839^41^, which acts upstream of GDH1, also sensitized Huh7 cells to JQ-1 treatment (Extended Data Fig.4a-c). Taken together, these results support the conclusion that BET inhibition induces GDH1 dependence in liver cancer cells, and GDH1 enzymatic activity is required for the viability of liver cancer cells upon BET inhibition.

### BET inhibition induces GDH1 expression through *miR-30a* downregulation

We next addressed how BET inhibition upregulates GDH1 expression in liver cancer cells. MYC, a master regulator of cellular metabolism, is the best-known BET target and mediates several oncogenic functions of BETs in cancer. Actually, targeting BET proteins has been considered as an actionable way to decrease MYC expression^27, 42^. We noted that BETi decreased MYC levels in liver cancer cells, and MYC targets were identified as top-ranked gene sets with altered expression by BETi (Fig. 2d and Extended Data Fig. 5a). We thus asked whether the functional effects of BET inhibition are associated with MYC expression. Unexpectedly, Huh7 cells with MYC overexpression were equally sensitive to JQ-1 or ABBV075, as both BETi comparably decreased cell growth in parental and MYC-overexpressed cells (Fig. 5a and Extended Data Fig. 5b). Different from BETi, MYC knock down didn’t increase but decreased GDH1 expression, while other targets (GLUT1, HK2 and LDHA) were downregulated similarly to BET inhibition (Fig. 5b). Growth of HepaMP9-1 cells, a murine HCC cell line driven by constitutive MYC overexpression and p53 loss (*MYC^OE^; Trp53^KO^*), was also impaired by dose-dependent JQ-1 or ABBV075 treatment (Extended Data Fig. 5c). GDH1 was consistently upregulated in HepaMP 9-1 cells treated by ABBV075 or ARV-825, a BRD4 proteolysis-targeting chimera (PROTAC) (Fig. 5c). Importantly, neither ABBV075 nor ARV-825 altered MYC expression in HepaMP9-1 cells, collectively suggesting that BET inhibition upregulates GDH1 expression largely through MYC-independent mechanisms.

**Fig. 5.**
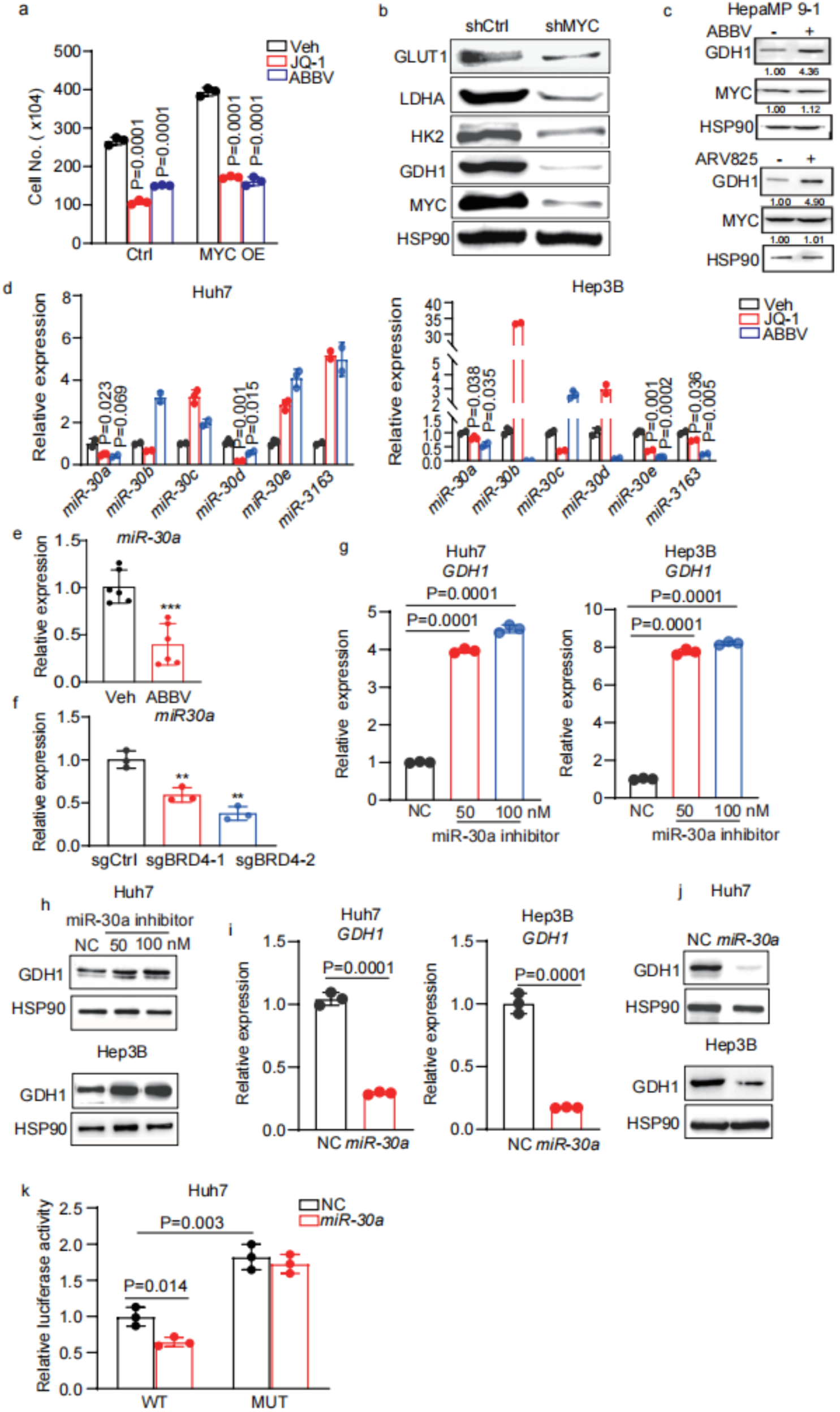
BET inhibition induces *GDH1* expression through *miR-30a* downregulation. **a**, Quantification of cell number in control and MYC-overexpressed Huh7 cells treated with BETi. **b**, Western blot analysis of indicated proteins on control and MYC knockdown Huh7 cells. HSP90 is used as a loading control. **c,** Western blot analysis of GDH1 and MYC expression in HepaMP cells treated with ABBV (5 nM) and ARV825 (5 nM) for 48 h. HSP90 serves as a loading control. **d**, Q-PCR analysis of indicated miR30 family genes in vehicle and BETi-treated Huh7 and Hep3B cells. **e**, Q-PCR analysis of *miR30a* levels in vehicle (n=5) and ABBV075 (n=6) treated Huh7 xenografts. **f,** Q-PCR analysis of *miR30a* expression from sgCtrl and sgBRD4 Huh7 cell lysates.**g,** Q-PCR analysis of *GDH1* mRNA levels in Huh7 and Hep3B cells transfected with control and inhibitor at indicated doses.**h**, Western blot analysis of *GDH1* expression in Huh7 and Hep3B cells transfected with control and inhibitor at indicated doses. **i**, Q-PCR analysis of *GDH1* mRNA levels in control and miR-30a-overexpressed Huh7 and Hep3B cells. **j,** Western blot analysis of *GDH1* expression in control and miR-30a-overexpressed Huh7 and Hep3B cells. **k**, Luciferase assay in Huh7 cells transfected with indicated reporter, control and expression plasmids.All statistical graphs show the mean ± s.e.m. *P* values were calculated using a two-tailed Student’s t-test. All experiments were performed in biological triplicate.

Increased mRNA levels result from either transcriptional activation and/or post transcriptional regulation such as elevated mRNA stability. Since BET proteins generally facilitate gene transcription, one would predict that BETi will inhibit gene transcription. Associated with elevated *GDH1* mRNA levels, we found a nearly 2-fold increase in the mRNA stability after JQ-1 exposure (Extended Data Fig. 5d). Therefore, we explored post transcriptional regulation of *GDH1* by miRNAs, which negatively regulate target mRNA stability and gene expression, although we couldn’t exclude other potential transcriptional regulation mechanism. Using the online tool TargetScan to predict *GDH1*-bound miRNAs, the miR30 family was identified as the only one potentially targeting *GDH1* 3’ untranslated region (UTR), which was conserved across several species. Among the five family members (miR- 30a-e), only miR-30a-5p (miR-30a) has consistent 30-50% decrease in expression in Huh7 and Hep3B cells by JQ-1 or ABBV075 treatment (Fig. 5d). Similarly, ABBV075 treatment reduced miR-30a levels in Huh7 xenografts (Fig. 5e), and BRD4 deficiency also decreased miR-30a levels in Huh7 cells (Fig. 5f). We thus hypothesized that miR-30a is the major miRNA targeting *GDH1.* To validate this, we performed both inhibition and over expression experiments and evaluated the effects on GDH1 expression. Transfection of inhibitor interfering endogenous miR-30a’s action increased both mRNA and protein levels of GDH1 in Huh7 and Hep3B cells (Fig. 5g, h). Conversely, ectopic miR-30a expression decreased *GDH1* mRNA and protein levels in both lines (Fig. 5i,j and Extended Data Fig. 5e). We further mutated the miR-30a target sequence and fused wild type (WT) or mutant (MUT) *GDH1* 3’UTR to C-terminal of Luciferase coding sequence (Extended Data Fig. 5f). Through dual luciferase assay, co-transfection of miR-30a decreased the relative activity of Luciferase-WT UTR reporter compared to vector control group, while such effects were abrogated in cells transfected with Luciferase-MUT UTR reporter vector (Fig.5k). These results provided substantial evidence to support the notion that miR-30a binds to 3’UTR and negatively regulates *GDH1* mRNA stability. Analysis of database finally further revealed a significant negative correlation between miR-30a and *GDH1* transcript levels (Extended Data Fig. 5g). In summary, we concluded that BET inhibition reduces miR- 30a levels which in turn increases *GDH1* mRNA stability leading to elevated protein levels.

### BET inhibition is synthetic lethal to OXPHOS blockage in liver cancer

Because BETi-treated liver cancer cells are able to maintain energy homeostasis involving enhanced glutamine oxidative metabolism and anaplerosis for TCA cycle, we finally hypothesized that targeting oxidative phosphorylation (OXPHOS) would disrupt the energy balance and sensitize the cancer cells to BET inhibition. To address this, we used a potent OXPHOS inhibitor IACS010759 (IACS)^43^ to treat cancer cells and compared growth among different groups. Compared to vehicle treatment group, either JQ-1 or IACS single treatment effectively slowed down Huh7 and Hep3B cell growth without obvious cell death, and combined JQ-1/IACS exposure more dramatically prevented cell growth and induced significant cell death (Fig. 6a, b and Extended Data Fig. 6a). JQ-1 and IACS had a strong synergistic effect, which was also evident between ABBV075 and IACS (Fig. 6c and Extended Data Fig. 6b). Combined JQ-1/ IACS or ABBV/IACS exposure significantly induced apoptosis in Huh7 cells, based on increased cleaved Caspase 3 levels (Extended Data Fig. 6c). Importantly, in the mouse liver cancer cell line HepaMP9-1 driven by MYC overexpression and p53 loss, JQ-1/IACS co-exposure also significantly decreased the cell viability (Extended Data Fig.6d). However, both SNU449 cells, a liver cancer cell line much less sensitive to JQ-1 than Huh7 and Hep3B cells, and IMR90, a non-cancer cell line, didn’t respond to combined JQ-1/IACS exposure, as evidenced by the comparable cell number and viability among different treatment groups (Extended Data Fig. 6e,f). These results indicate that only BETi-responsive liver cancer cells are more sensitive to JQ-1/IACS combination therapy.

**Fig. 6.**
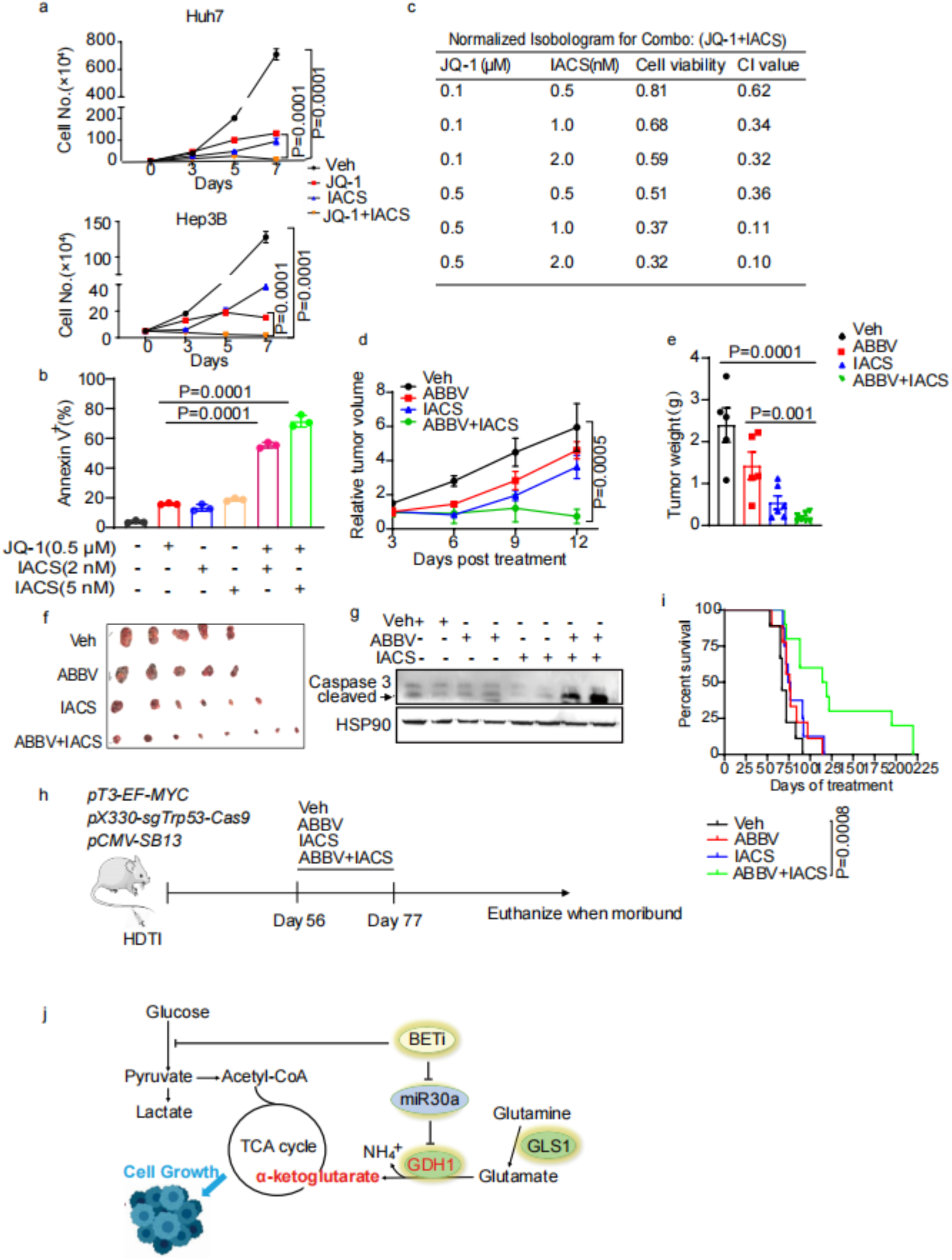
BET inhibition is synthetic lethal to OXPHOS blockage in liver cancer. **a**, Huh7 and Hep3B cell growth assays from the indicated treatment groups with JQ1 and IACS. **b**, Statistic analysis of the percentage of annexin V^+^ Huh7 cells of indicated treatment groups for 6 days. **c**, Synergistic effects between JQ-1 and IACS treatment of Huh7 cells. CI values<1 indicates synergy between two inhibitors. **d**, Quantification of Huh7 xenograft tumor growth in vehicle (n=5), ABBV (n=5), IACS (n=5) and ABBV/IACS combination (n=8) treatment cohorts. **e**, Quantification of Huh7 xenograft tumor weights in vehicle (n=5), ABBV (n=5), IACS (n=5) and ABBV/IACS combination (n=8) treatment cohorts. **f** Representative Huh7 xenograft tumor images in vehicle, ABBV075, IACS and ABBV/IACS combination treatment cohorts. **g**, Western blot analysis of cleaved caspase 3 in Huh7 xenograft tumor combined ABBV and IACS exposure. **h**, A scheme of liver tumorigenesis and drug treatment of *MYC^OE^; Trp53^ko^* mouse model. **i**, Kaplan–Meier survival curves of vehicle (n=8), ABBV075 (n=8), IACS (n=9) and ABBV/IACS combination (n=10) treatment cohorts. A log-rank Mantel–Cox test was performed between the groups. All statistical graphs except (**i**) show the mean ± s.e.m. Except (**i**)*, P* values were calculated using a two-tailed Student’s t-test. All experiments were performed in biological triplicate.

To validate BETi/OXPHOSi combination therapy potential *in vivo*, we first performed treatment on Huh7 xenograft tumor model using ABBV075 and IACS. As shown in Fig. 6d, compared to vehicle cohort, either 12-day ABBV075 or IACS monotherapy substantially blunted tumor growth, and IACS seemed to have a stronger inhibitory effect. Notably, ABBV075/IACS combination therapy more potently blunted tumor growth, and the endpoint tumor weights were much smaller than any other cohorts (Fig. 6e,f). Consistent with in vitro treatment, xenografts exposed to combined ABBV and IACS exhibited much higher cleaved caspase 3 level than other groups (Fig. 6g).

We then performed drug treatment on a genetically engineered mouse model driven by MYC overexpression and p53 loss as previously^44^. In this model, a MYC-expressing transposon vector and another expressing Cas9 and *Trp53* sgRNA were delivered to murine hepatocytes using hydrodynamic tail vein injection, where MYC overexpression and p53 loss trigger rapid liver tumorigenesis and shorten animal survival. After injection, mice randomized in different groups received three-week mono or combination therapy and their survival was monitored (Fig. 6h). Mice receiving either ABBV075 or IACS monotherapy exhibited longer survival than vehicle controls. Most importantly, the longest overall survival was observed with combined ABBV075 and IACS administration (Fig.6i). Notably, wild type mice receiving combined ABBV075 and IACS treatment initially lost about 10% body weight but progressively recovered after treatment, and the liver histology between two cohorts were indistinguishable by pathological examination (Extended Data Fig. 6g,h). Based on above results, we conclude that BETi combined with OXPHOS inhibitor represents an enhanced therapeutic option in liver cancer.

## Discussion

BET proteins regulate multiple gene expressions involved in cancer pathogenesis and are emerging therapeutic targets. Results from the first clinical trials confirm the antitumor potential of BET inhibitors (BETi), but their efficacy as single agents seem to be limited. Instead, combination therapies and next generation of compounds such as BET degraders may open new possibilities for targeting BET proteins in cancer^25, 26^. Ongoing BETi-based combination therapies include kinase inhibitors, immune modulators, epigenetic drugs and hormone therapy, and the clinical outcomes are awaited. Notably, expression of several metabolic genes can be epigenetically regulated by BET proteins, and BET target genes including MYC are master regulators of cellular metabolism. It is thus reasonable to postulate that targeting BET proteins may directly or indirectly affect cancer metabolism, rendering BETi-treated cells dynamically undergo metabolic remodeling in this context. In support of this notion, HDAC inhibition elicits metabolic reprogramming by targeting super-enhancers in glioblastoma, and combination treatment with HDAC and FAO inhibitors exhibits a better therapeutic potential^45^. However, how BETi induces metabolic remodeling and whether a metabolic targeting strategy can be combined with BETi for cancer treatment has not been explored. Here using liver cancer as a model, we uncover a GDH1-dependent glutamine metabolic remodeling upon BET inhibition, and provide the a proof-of-principle for “synthetic lethal” targeting GDH1-OXPHOS metabolic axis and BET proteins for improved cancer treatment (Fig. 6j).

We find that both BRD2 and BRD4 but not BRD3 are required for liver cancer cell growth, and even transient BET suppression using pan-BET inhibitors (JQ-1 and ABBV075) is sufficient to impede cell proliferation and xenograft tumor growth. From a therapeutic perspective, short-term treatment may also reduce the side effects of BET inhibition. To address how BET inhibition may affect liver cancer cell metabolism, we start from RNA-sequencing and find that BETi decreases expressions of genes in glucose and mitochondrial metabolism, consistent with the notion that BET proteins generally facilitate gene transcriptional activation. Interestingly, GDH1 is identified as one of few metabolic genes upregulated by BET inhibition. Through the production of α-KG, GDH1 contributes to redox homeostasis in breast and lung cancer cells^40^, promotes anoikis resistance in LKB1-deficient lung cancer^46^, and is required for glioblastoma cells to survive under glucose limitation^47^. In line with GDH1 upregulation, BETi increases not only the abundance of multiple glutamate metabolism and TCA cycle metabolites, but also the activity of glutamine oxidative metabolism, which provides anaplerotic metabolites including α-KG for TCA cycle. As a result, BETi-treated cells are able to maintain energy homeostasis despite the impaired growth. Functionally, targeting GDH1, GLS1 or OXPHOS consistently results in synthetic lethality with BETi, while none of these single treatments induces dramatic cell death. Collectively, these findings demonstrate that BETi-treated cancer cells undergo GDH1-mediated glutamine metabolic remodeling to maintain energy homeostasis and viability, blocking this adaptation process elicits energy crisis and synthetic lethality with BET inhibition.

Context-dependent GDH1 modulation can occur at different levels. For example, the transcription factor PLAG1 controls GDH1 expression during anoikis resistance^46^, the protein but not mRNA level of GDH1 is induced when proliferating mammary epithelial cells undergo quiescence^48^, and GDH1 enzymatic activity but not protein level is elevated when glucose oxidation is inhibited in glioblastoma cells^47^. We identify a distinct mechanism of GDH1 upregulation by BET inhibition in liver cancer cells, which is through downregulation of miR-30a independent of MYC. We demonstrate that miR-30a negatively regulates *GDH1* mRNA stability, and miR-30a levels are reduced upon BET inhibition. Although miR-30a downregulation may affect multiple gene expressions, our results support that GDH1 is the one involved in glutamine medtabolic remodeling upon BET inhibition. It remains unclear how BETi decreases miR-30a levels and whether other mechanisms also regulate GDH1 in this context.

Finally, we show that simultaneously targeting BET proteins and GLS1-GDH1-OXPHOS metabolic axis limits liver tumor growth, even in tumors driven by ectopic MYC overexpression. In our MYC/p53 model, three-week ABBV075/IACS combination therapy significantly extends the survival of animals compared to monotherapy cohorts. This treatment regimen seems to be well-tolerated, despite that the treated animals initially lose and progressively recover the body weight. It should be noted that recent clinical trial reveals that IACS monotherapy is associated with toxic side effects including peripheral and optic neuropathy, and hence only modest evidence of enhanced efficacy is observed in patients with acute myeloid leukemia or advanced breast cancer and pancreatic cancer^49^. This observation strongly suggests that it’s necessary to either identify the cancer type/patients suitable for the treatment, or optimize the treatment regimen to reduce the side effects of targeting complex I. Indeed, complex I loss exposes fermentation as a therapeutic target in hurthle cell carcinoma and has implications for other tumors bearing mutations that irreversibly damage mitochondrial respiration^50, 51^. We herein show that BET inhibition on the one hand has a strong negative impact on glycolysis, on another hand increases GDH1-mediated glutamine metabolic remodeling, establishing a synthetic lethality to target both pathways. Notably, BETi/OXPHOSi combination therapy is more effective to BETi-responsive liver cancer cells, as non-cancerous cells (IMR90) and BETi-less sensitive SNU449 cancer cells don’t respond to this treatment, raising a remaining question about how to stratify liver cancer patients that would benefit from the combination therapy. In addition to OXPHOS inhibition, our findings also suggest that targeting induced metabolic dependency upstream of OXPHOS, such as GLS1 or GDH1, can also induce synthetic lethality to BETi-responsive liver cancer. As new generation of BET inhibitors are emerging, such as BRD4 PROTAC (ARV-825), together with suitability of GLS1 and GDH1 as drug targets, one can anticipate that targeting BET proteins and GLS1-GDH1 metabolic axis may hold therapeutic potential, which is worth to be fully evaluated in the future.

In summary, our study provides mechanistic insights into the metabolic adaptation underlying BET inhibition in liver cancer, and proposes an option for synthetic lethal targeting GDH1-dependent glutamine metabolism and BET proteins for improved combination treatment of liver cancer.

## ACKNOWLEDGEMENTS

We thank the members of the Li laboratories for their helpful discussions and insights on the manuscript. We are grateful to Jinyang Li for help with processing the RNA-sequencing data. We are also grateful to the core facility of metabolomics for technical assistance. This work was supported by the National Key R&D Program of China (2022YFA1103900 to F.Li), the National Natural Science Foundation of China (82273223 to F.Li, 32270798 to P. Liu).

## AUTHOR CONTRIBUTIONS

W.M. and F.L. conceived the project and designed the experiments, P.L. and F.L. supervised the overall study. W.M., L. L., N.A., N.L., M.B., J.P. and A. M. performed the experiments. W.M. and F.L. analyzed the data and wrote the manuscript. All authors revised and approved the manuscript.

## METHODS

### Cell culture

Huh7, Hep3B, IMR90, SNU449, 293T cell lines were purchased from ATCC. HepaMP-9-1 cell line was generated in the Simon lab at the University of Pennsylvania. Huh7, Hep3B, 293T and HepaMP9-1 were cultured in DMEM medium containing 10% fetal bovine serum (FBS) (hereafter growth medium). SNU449 cells were cultured in RPMI medium containing 10% FBS. The cells were routinely tested to exclude mycoplasma contamination. For cell growth assays, single cells in complete medium were seeded in 6-well plates (5×10^4^/well, in triplicate), and the growth medium was changed every 2 days, cell cumbers were counted with a hemacytometer. For clonogenicity assay, single cells were seeded in 12-well plates (2×10^3^/well, in triplicate) and cultured for 10-12 days, the medium was changed every 3 days. Fur suspension culture, single cells were seeded on ultra-low attachment culture dishes (Corning) in DMEM/F12 serum-free medium (Invitrogen)^29^, which contained 2 mM L-glutamine, 1% sodium pyruvate (Invitrogen), 100 μg/ml penicillin G, and 100 U/ml streptomycin supplemented with 20 ng/ml epithelial growth factor (Invitrogen), 10 ng/ml fibroblast growth factor-2 (Invitrogen), N2 (Invitrogen), and B27 (Invitrogen). The cells were incubated in a CO2 incubator for two weeks, and numbers of oncosphere cells were counted under a stereomicroscope (Olympus, Tokyo, Japan).

### Crystal violet staining

The cultured cells were washed once with 1xPBS, then stained with 1xPBS solution containing 20% methanol and 0.25% crystal violet for 30 min, the cells were then washed to remove residual crystal violet.

### Chemicals

JQ-1 (#A1910), CB839 (#B4799) were from APExBIO; ABBV075 (#HY-100015), ARV-825 (#HY16954), IACS010759 (#HY-112037) were from MedChemExpress; R162 (#E1170) was from Selleck; [U-13C]-glutamine and [U-13C]-glucose were from Cambridge Isotope Laboratories; acetonitrile (ACN), methanol (MeOH), formic acid at liquid chromatography grade were from Thermo Fisher company; methylene diphosphate was from Sigma.

### Western blot analysis

Cells were washed with 1x PBS and lysed using lysis buffer (150 mM NaCl, 10 mM Tris pH 7.6, 0.1% SDS and 5 mM EDTA) containing Halt Protease and Phosphatase Inhibitor Cocktail (Thermo Fisher Scientific, 78445). Samples were centrifuged at 12,000 rpm for 20 min at 4°C. Protein lysates were separated by SDS-PAGE and were transferred to PVDF membranes (Millipore). All membranes were incubated with the indicated primary antibodies diluted in TBS-T (20mM Tris pH 7.5, 150 mM NaCl, 0.1% Tween-20) with 5% bovine serum albumin (Sigma-Aldrich, A7906) overnight at 4 °C. After TBS-T washes, membranes were incubated with secondary antibody and developed with Western Lightning Plus-ECL, Enhanced Chemiluminescence Substrate (PerkinElmer, cat. NEL103E001EA). The following primary antibodies were used: Cell signaling technology: Phospho-AMPKα (Thr172) (40H9) Rabbit mAb (#2535), AMPKα (#2532); Abclonal:HSP90 (#A5027), HK2 (#A0994), GLUT1 (#A11208), LDHA (#A0861), GDH1(#A5176), GLS (#A11043); Proteintech: BRD4 (#67374), MYC (#67447-1-Ig), GOT1 (#14886-1-AP), GOT2 (#14800-1-AP), GPT1 (#16897-1-AP), GPT2(#16757-1-AP), PSAT1(#10501-1-AP), TFAM (#23996-1-AP); Santa Cruz: BRD2 (#sc-514103), BRD3 (#sc-81202).

### Reverse transcription (RT) and Quantitative real-time PCR (Q-PCR)

Total RNA was isolated using RNeasy Mini Kit (Qiagen, Cat. 74104). cDNA was synthesized using a High-Capacity RNA-to-cDNA kit (Vazyme, Cat. R323-01). Q-PCR was performed using ViiA7 Real-Time PCR system (Applied Biosystems) with SYBR green master mix (Vazyme, Cat. Q711-02). The relative mRNA levels were normalized to 18S ribosomal RNA. Q-PCR primers for miRNAs were summarized in the supplementary Table 1. Other Q-PCR primers were summarized in the supplementary Table 2.

### PI/Annexin V staining and flow cytometry

Cell death was determined using the FITC–Annexin V, PI Kit (#556547) from BD Biosciences according to the manufacturer’s instructions. Flow cytometry was performed using the BD FACS Calibur instrument (BD FACS Calibur), with dead cells represented as Annexin V-positive population, resultant data were analyzed with the FlowJo 10.6.2 software (https://www.flowjo.com).

### Metabolomics

Huh7 cells were treated with vehicle or JQ-1 (0.5 μM) for 48 hours, and then replated in 10 cm plates overnight. For metabolite extraction, medium was removed by aspiration and metabolism was immediately quenched (without any washing steps) by adding 500 μl −20 °C 40:40:20 acetonitrile: methanol: water containing 0.5% formic acid. After about 5∼10 s, cell extract was neutralized with 44 μl 15% ammonium bicarbonate. The cell extracts were transferred to 1.5 ml tubes and stored at −80 °C overnight and further cleared by centrifuging to remove protein at 15,000 × g for 10 minutes. The supernatant was used for LC-MS analysis. Targeted metabolomics analysis was performed using Shimadzu Prominence HPLC system (ExionLC AD) interfaced with a QTRAP 6500+ system (AB SCIEX). The sample injection volume was 5 µl. Metabolites were separated through an iHILIC-(P) Classic HILIC column (100 × 4.6 mm with 3.5 μm particle size, Waters) with column temperature maintained at 40 °C. Mobile phases consisted of 20 mM ammonium acetate in 25 mM ammonia water (mobile phase A) and acetonitrile (mobile phase B) and was run at a flow rate of 0.4 ml/min. The gradient was as follows: 0 min, 85% B; 0.1 min, 85% B; 3.5 min, 32% B; 12 min, 2% B; 16.5 min, 2% B; 17 min, 85% B; 26 min, 85% B. The mass spectrometer was run in multiple reactions monitoring (MRM) mode. The ESI source parameters: source temperature of 475 °C, the ion source gas 1 and 2 at 60 psi, the curtain gas at 35 psi, the ion spray voltage at 4850 V or −4500 V for positive or negative modes. Data were processed and analyzed using SCIEX OS software.

### [U-13C]-glucose and [U-13C]-glutamine labeling and LC-MS

All ^13^C studies were performed in DMEM medium containing 10% dialysed FBS and prepared so that 100% of either the glucose or glutamine pool was labelled with ^13^C, and the other pool was unlabelled. Huh-7 cells were treated with vehicle or JQ-1 (0.5 μM) for 48 hours, and replated in 6 well plates (10^6^/well) overnight, then the cells were switched to fresh culture medium so that cells should reach 60-90% confluence next day. At the beginning of the experiment, cells were switched into labeled [U-13C]-glucose or [U-13C]-glutamine DMEM medium containing 10% dialyzed FBS and incubated for 4 h. The metabolite extraction and LC-MS sample preparation steps were identical to targeted metabolomics experiment. Isotope labeling data acquisition was performed using a Shimadzu LC system coupled to a TripleTOF mass spectrometer (QTOF 6600+, ABSciex, made in Woodlands Central Industrial Estate, Singapore). Chromatographic separation was achieved using a HILIC column (IHILIC-(P) Classic column, 5 μm, 150 x 2.1 mm, 200 A, made in Sweden). The mobile phase A was 20 mM ammonium acetate, 0.1% ammonium hydroxide, and 2.5 μM methylene diphosphate in 95: 5 water/ACN and mobile phase B was ACN. The gradient was as follows: 0 min, 85% B; 2 min, 85% B; 7 min, 65% B; 12 min, 35% B; 12.1 min, 20% B; 15.9 min, 20% B; 16 min, 85% B; 23 min, 85% B. The flow rate was set at 0.2 mL/min with a sample injection volume of 5 µl and the total run time at 23 min. ESI parameters setup: GS1, 60; GS2, 60; CUR,35; temperature, 500; ISVF, −4500 in negative modes. For each metabolite, the area of ^12^C and various ^13^C isotopologue peaks were integrated using El-MAVEN software (version 0.12.1). Correction for natural isotope abundance was performed using R package AccuCor. The labeling fraction of a specific isotopologue was calculated as the area of that isotopologue divided by the sum of all isotopologues’ area^52^.

### Constructs and cell transfections

shRNAs and were cloned into pLko.1-puro (Addgene #8453) linearized with AgeI and EcoRI. sgRNAs were cloned into pLentiCRISPR v2 Lko.1-puro (Addgene #52961) linearized with BsmBI. The following oligonucleotides for shRNAs were used: *shBRD2* CCCTGCCTACAGGTTATGATT; *shBRD3* GCTGATGTTCTCGAATTGCTA; *shBRD4* CCTGGAGATGACATAGTCTTA; *shGDH1-1* GCCATTGAGAAAGTCTTCAAA; *shGDH1-2* GCAGAGTTCCAAGACAGGATA; *shMYC* CCTGAGACAGATCAGCAACAA. The following oligonucleotides for sgRNAs were used: *sgBRD4-1*: TAAGATCATTAAAACGCCTA, *sgBRD4-2*: TCTTCCTCCGACTCATACGT. To produce lentiviruses, 293T cells were co-transfected with LentiCRISPR v2 plasmids along with packaging plasmids psPAX2 and pMD2.G using PEI transfection reagent (#23966-100, Polysciences). Lentiviruses were collected 48 hours after transfection. MiR-30a inhibitor was purchased from GenePharma, and transfected using Lipofectamine™ RNAiMAX Transfection Reagent (ThermoFisher Scientific, #13778075) following the manufacturer’s instructions. For luciferase reporter assays, *GDH1* 3’UTR was amplified using the following primers (Forward: ATCGCTCGAGTGATGAAAGCTGCGCACTAGTTCTGCAGACCTATCACAAGT; Reverse: ATCGAAGCTTAAGACTATGCTTTCAGGGAT) and cloned into pMIR-REPORT™ miRNA expression reporter vector. Mutant 3’UTR reporter was generated through site-directed mutagenesis and confirmed by DNA sequencing. Luciferase activity was quantified using a dual luciferase assay kit (Promega). Relative luciferase activity was calculated by normalizing relative luciferase activities to Renilla activities in each well.

### Animal experiments

All mouse experiments were reviewed and approved by the Institutional Animal Care and Use Committee at Fudan University. Male C57BL/6, female NCG mice were purchased from GemPharmatech Co.,Ltd (Nanijng, China), and were maintained in a SPF animal facility at Fudan University. For subcutaneous injections, 5×10^6^ Huh7 cells mixed 1:1 with Matrigel (BD Biosciences, #356234) in a final volume of 100 μl were injected into the blanks of NCG mice, respectively. The tumor volume was monitored by caliper measurements. When tumor volume reached 100 mm^3^, NCG mice were randomized and divided into groups for drug treatment. For hydrodynamic tail vein injection (HDTI), vectors were prepared using the EndoFree-Maxi Kit (Qiagen) and resuspended in a sterile 0.9% NaCl solution/plasmid mix containing 10 μg of pT3-MYC (Addgene #92046), 10 μg of pX330-p53 (Addgene 59910), and 2.5 μg of CMV-SB13 transposase. A total volume mix corresponding to 10% of body weight was injected via lateral tail vein in 5–8 s into 6-week-old male C57Bl/6 mice, as previously. For drug treatment, ABBV075 and IACS010759 was diluted in 0.5% Methyl cellulose solution (MC), and administrated via oral gavage once daily at 1 mg/kg and 7.5 mg/kg, respectively. 0.5% Methyl cellulose solution was used as vehicles. The NCG mice were treated for 12 days, and HDTI-treated C57BL/6 mice were treated for 3 weeks. After treatment, the mice were carefully monitored, and euthanized when moribund, and tumor burden was confirmed. For HE staining, mouse tissues were fixed in 4% paraformaldehyde immediately after collection, dehydrated and stained with hematoxylin and eosin as previously^17^.

### RNA-seq and GSEA

Total RNA was extracted from vehicle and BETi-treated Huh7 cells using a RNeasy mini kit (Qiagen). The RNA quality test, library construction and sequencing were performed by Novogene Corporation (Beijing). For data analysis, Fastq files were checked for quality using FastQC and qualimap. Alignment was performed using the STAR aligner under default settings with the hg19 reference genome. Raw counts of gene transcripts were obtained from the resulting bam files using the feature Counts. The raw count matrix was subsequently imported into R-studio (R version 3.3.3) and used as input for DESeq2 following the vignette of the package for normalization and differential gene expression analysis. Salmon/Sailfish was used in parallel to normalize and quantitate gene expression in transcripts per million through quasi-alignment. Gene set enrichment analysis (http://software.broadinstitute.org/gsea/index.jsp) was run for the contrast in pre-ranked mode using the DESeq2 statistic as the ranking metric. Where there were redundant mappings, the statistic with the highest absolute value was chosen^53^.

### Statistics

All results were obtained from three independent biological experiments, using three technical replicates per condition, unless stated otherwise. Statistical analyses per experiment are indicated in figure legends. Statistical tests were performed in GraphPad Prism 9.0.0 using Student’s two-tailed unpaired t-test for pairwise comparisons, one-way analysis of variance (ANOVA) for multiple comparisons, two-way ANOVA for multiple comparisons involving two independent variables, or and by log-rank test for comparisons of survival distributions of two groups. Statistical data are presented as mean ± SEM of at least three independent experiments. A p-value less than 0.05 was considered significant.

## Data and Code Availability

The TCGA liver cancer dataset (https://cancergenome.nih.gov/cancersselected/LiverHepatocellularCarcinoma) was downloaded and analyzed at the Molecular Profiling Facility at the University of Pennsylvania as previously. All sequencing data have been deposited in the Gene Expression Omnibus under the series GSE184065. The software and algorithms for data analyses used in this study are all well-established from previous work and are referenced throughout the manuscript. No custom code was used in this study. All data supporting the findings of this study are available from the corresponding author upon reasonable request.

---

**Table 1.**
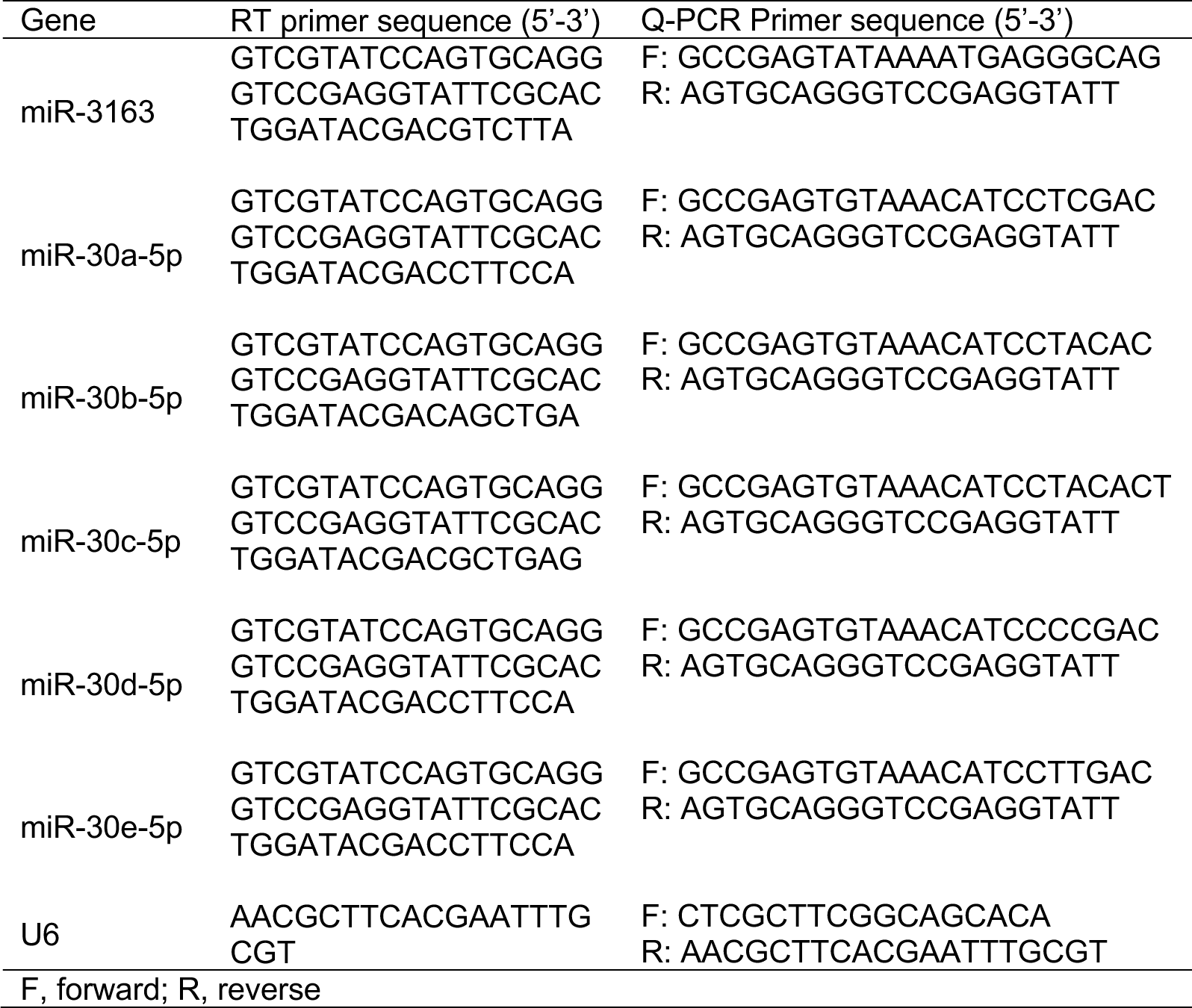
Primer sequences for microRNA reverse transcription (RT) and Q-PCR analysis.

**Table 2.**
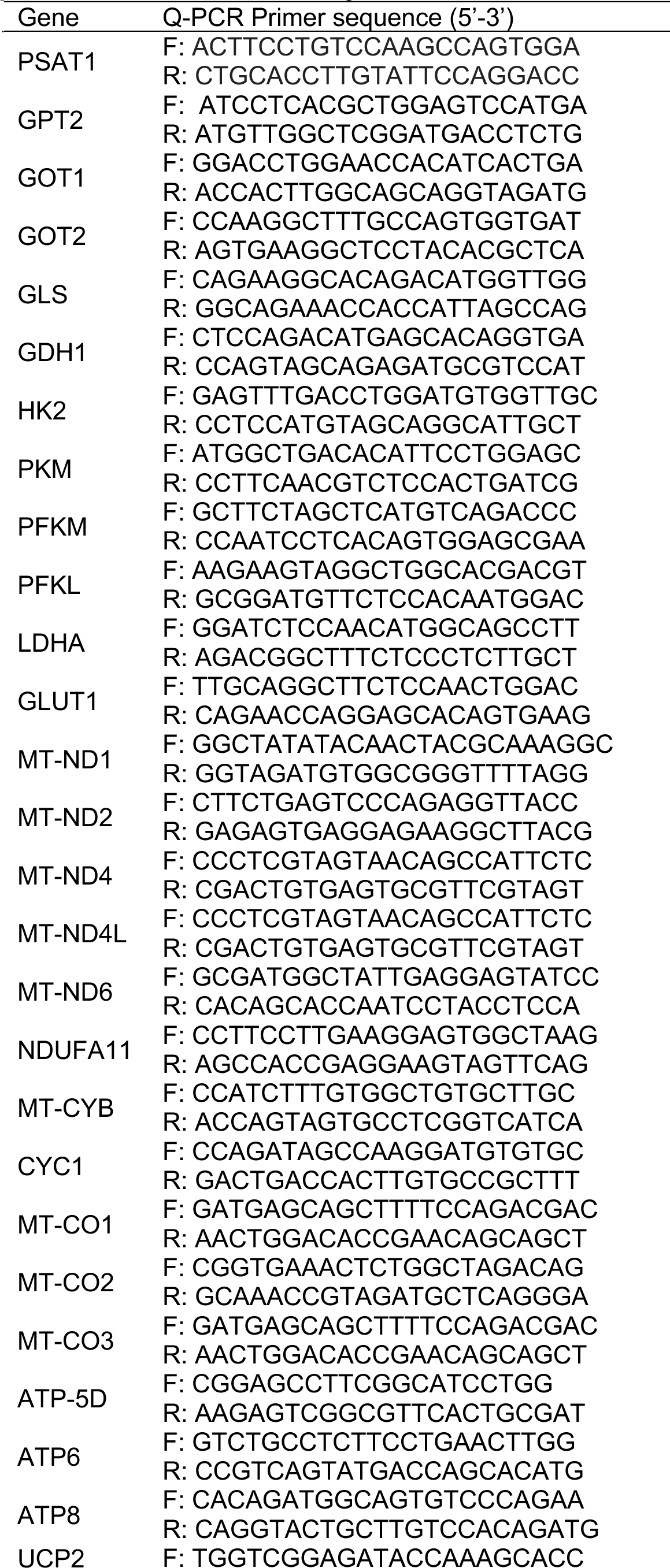
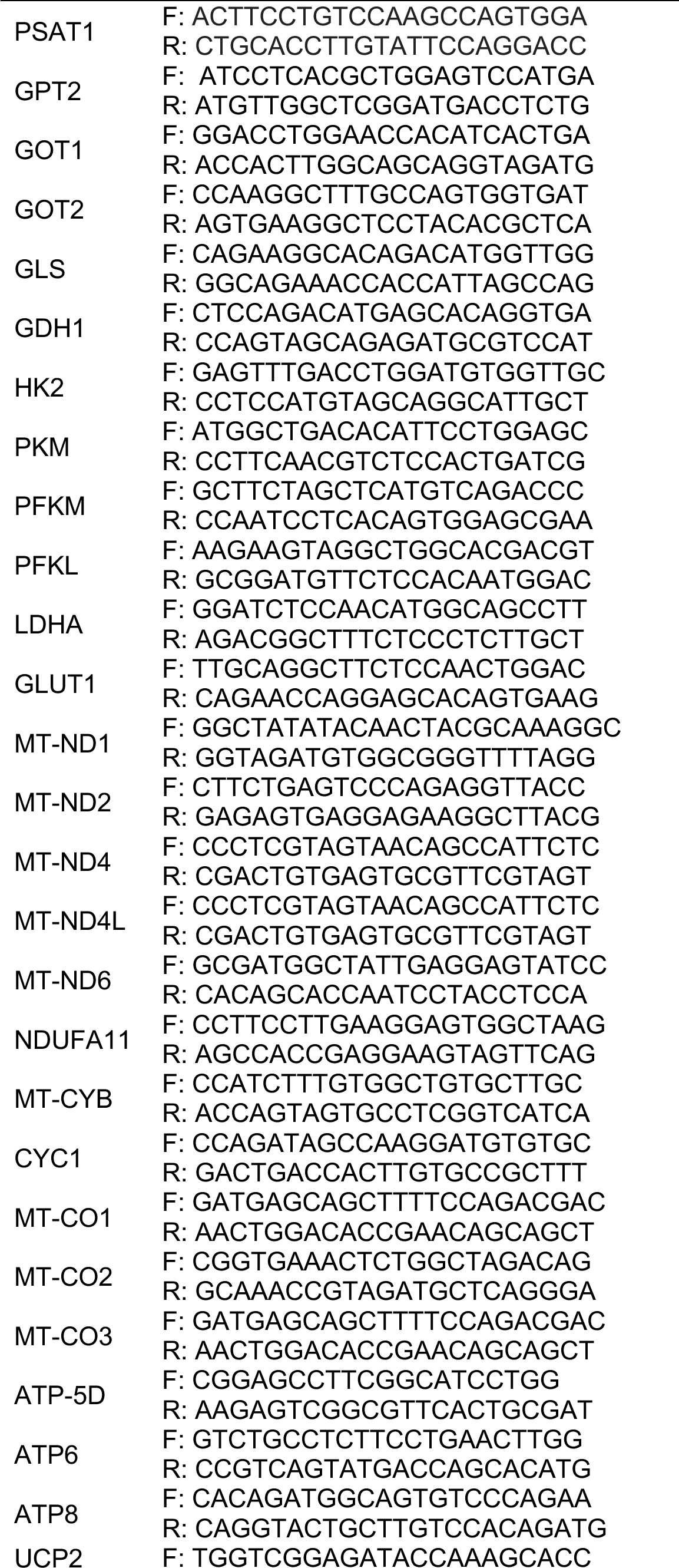
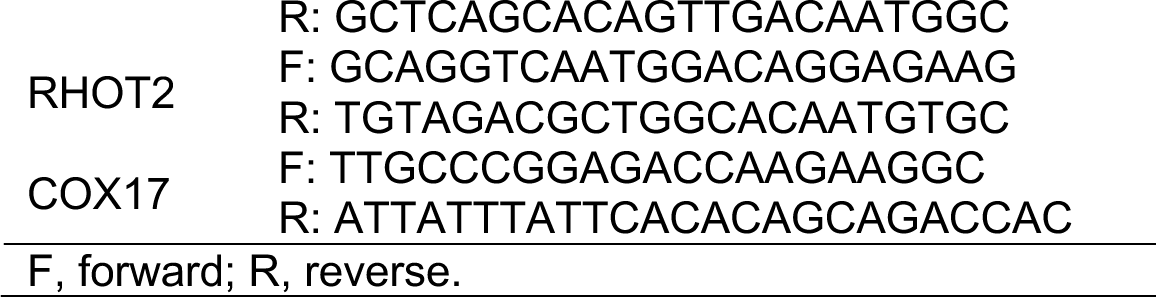
Primer sequences used for Q-PCR analysis.

