## Extended Data Figures and Legends for "BET inhibition induces GDH1-dependent glutamine metabolic remodeling and vulnerability in liver cancer"

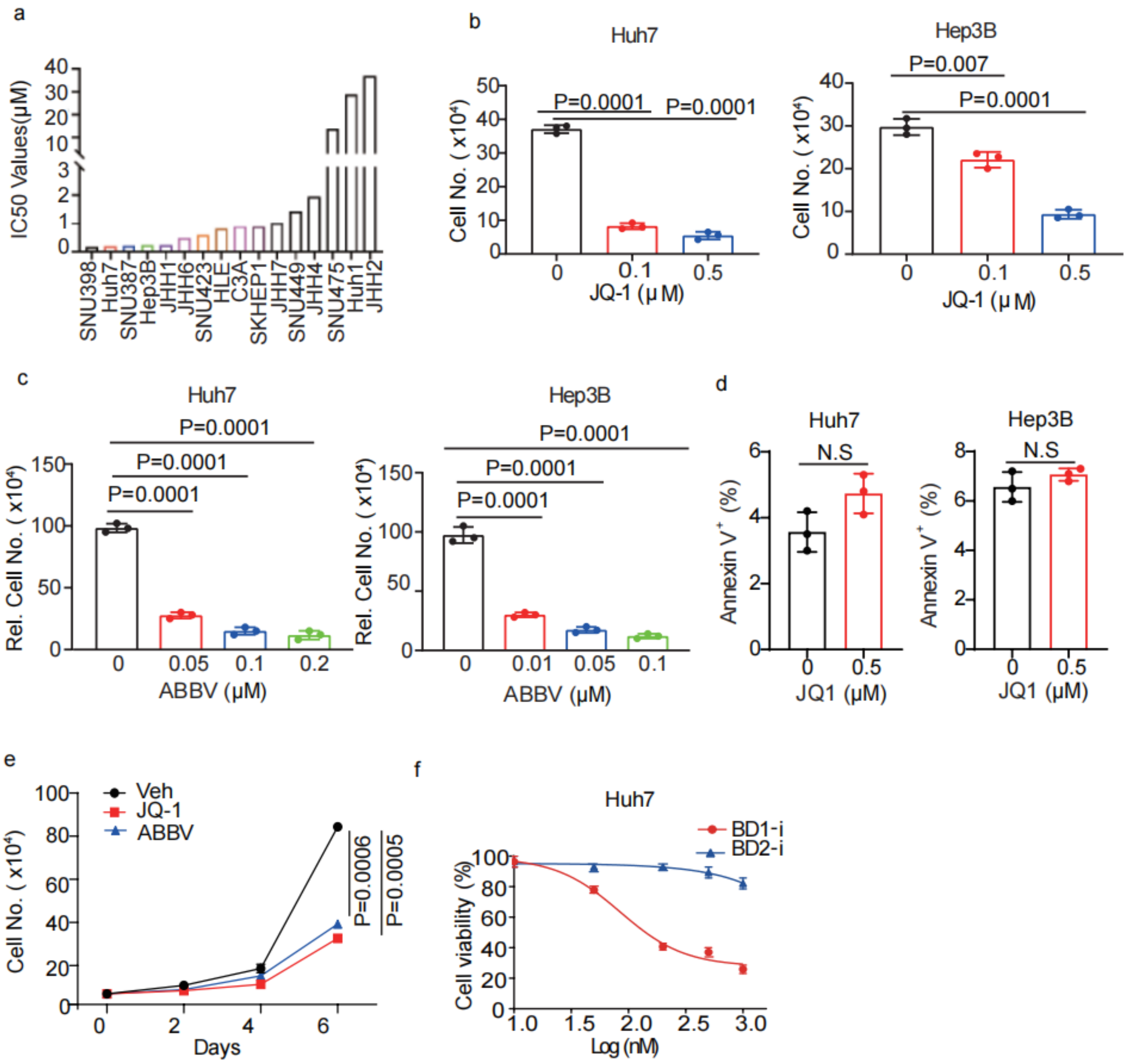


**Extended Data Fig. 1. BET inhibition blunts HCC cell grow and induces differentiation. a**, JQ-1 IC50 values across liver cancer cell lines from the "The Genomics of Drug Sensitivity in Cancer Project" **b**, Quantification of Huh7 and Hep3B cell numbers after JQ-1 treatment for 48 hours. **c**, Quantification of Huh7 and Hep3B cell numbers after ABBV-075 treatment for 48 hours. **d**, Statistic analysis of the percentage of annexin $V^+$ Huh7 and Hep3B cells in vehicle and JQ-1 treatment groups. **e**, Cell growth assays for replated Huh7 cells after vehicle, JQ-1 or ABBV-075 treatment for 48 hours. **f**, Relative cell viability in Huh7 cells treated with indicated doses of BD1 and BD2 inhibitors. All statistical graphs show the mean ± s.e.m. *P* values were calculated using a two-tailed Student's t-test. All experiments were performed in biological triplicate.

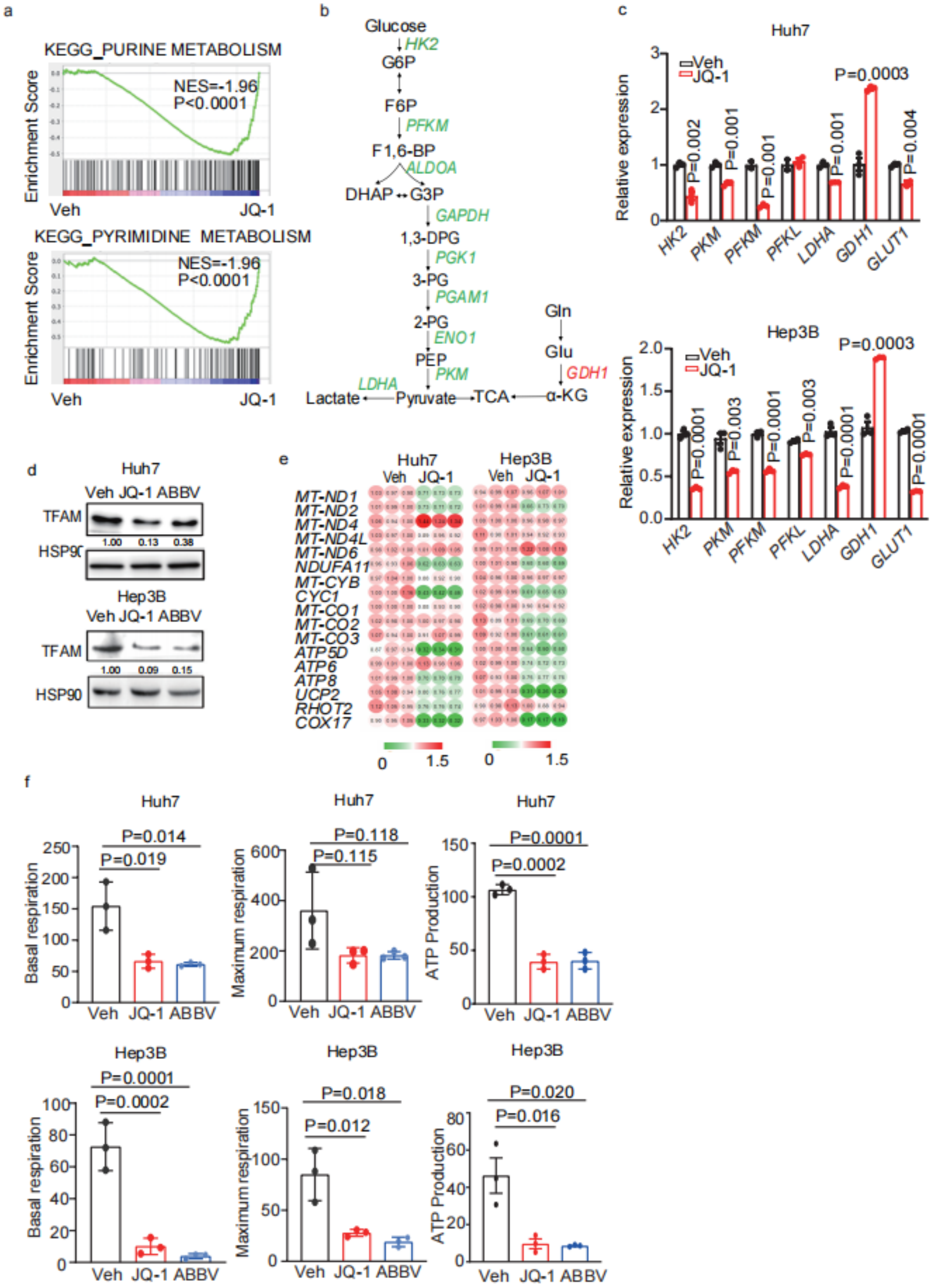


**Extended Data Fig. 2. Metabolic gene expression changes in liver cancer cells upon BET inhibition.**

**a**, GSEA for purine and pyrimidine metabolism from vehicle and JQ-1 groups based on RNA-sequencing data. **b**, A scheme of glycolytic gene expression change upon JQ-1 treatment of Huh7 cells. **c**, Q-PCR analysis of glycolysis gene expression in JQ-1-treated Huh7 and Hep3B cells. **d**, Western blot analysis of TFAM from vehicle and BETi-treated Huh7 and Hep3B cell lysates. HSP90 is used a loading control. **e**, Q-PCR analysis of mitochondrial ETC subunit gene expression in vehicle and JQ-1-treated Huh7 and Hep3B cells. **f**, Quantification of seahorse assays in vehicle and BETi-treated Huh7 and Hep3B cells. All statistical graphs show the mean ± s.e.m. *P* values were calculated using a two-tailed Student's t-test. All experiments were performed in biological triplicate.

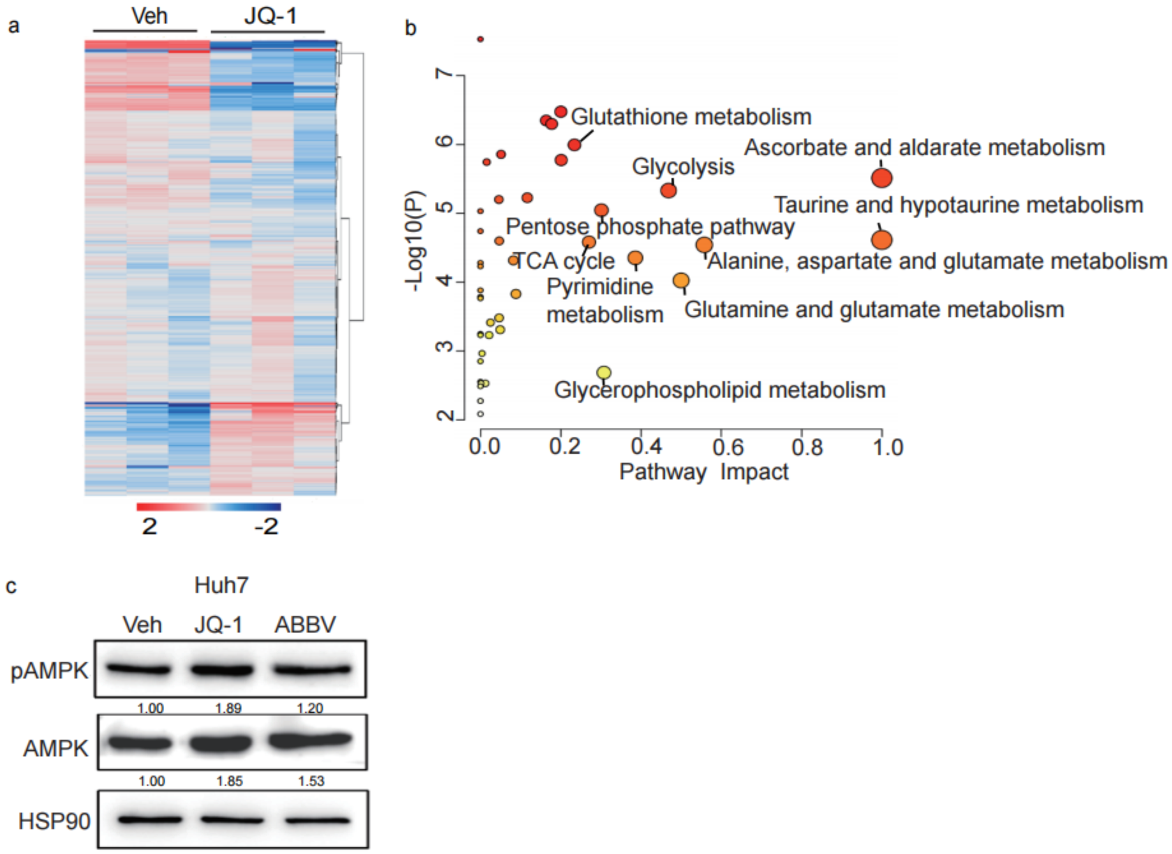


**Extended Data Fig. 3. Metabolomic profiling of liver cancer cells upon BET inhibition.** **a**, A heatmap of untargeted metabolomics in vehicle and JQ-1 treated Huh7 cells. **b**, Metabolic pathway analysis based on untargeted metabolomics in (**a**). **c**, Western blot analysis of AMPK and pAMPK levels in vehicle and BETi-treated Huh7 cell lysates. HSP90 is used a loading control. All statistical graphs show the mean ± s.e.m. *P* values were calculated using a two-tailed Student's t-test. All experiments were performed in biological triplicate.

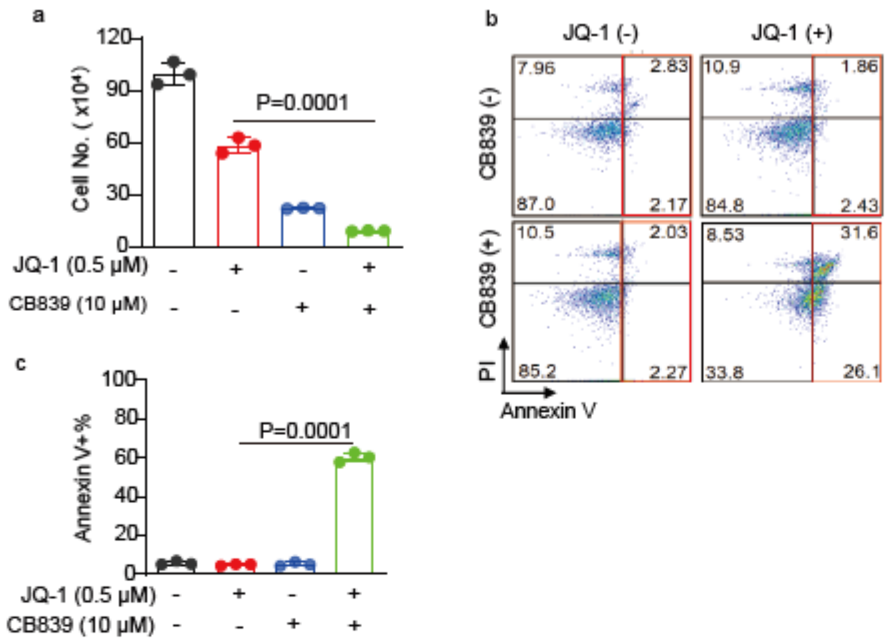


**Extended Data Fig. 4. Targeting GLS1 sensitizes liver cancer cells to BET inhibition. a**, Quantification of cell numbers for Huh7 cells treated with JQ-1 and/or CB839. **b,** Representative flow cytometry plot of PI and annexin V staining of Huh7 cells from indicated groups.**c**, Statistic analysis of the percentage of annexin $V^+$ Huh7 cells in the indicated treatment groups. All statistical graphs show the mean ± s.e.m. *P* values were calculated using a two-tailed Student's t-test. All experiments were performed in biological triplicate.

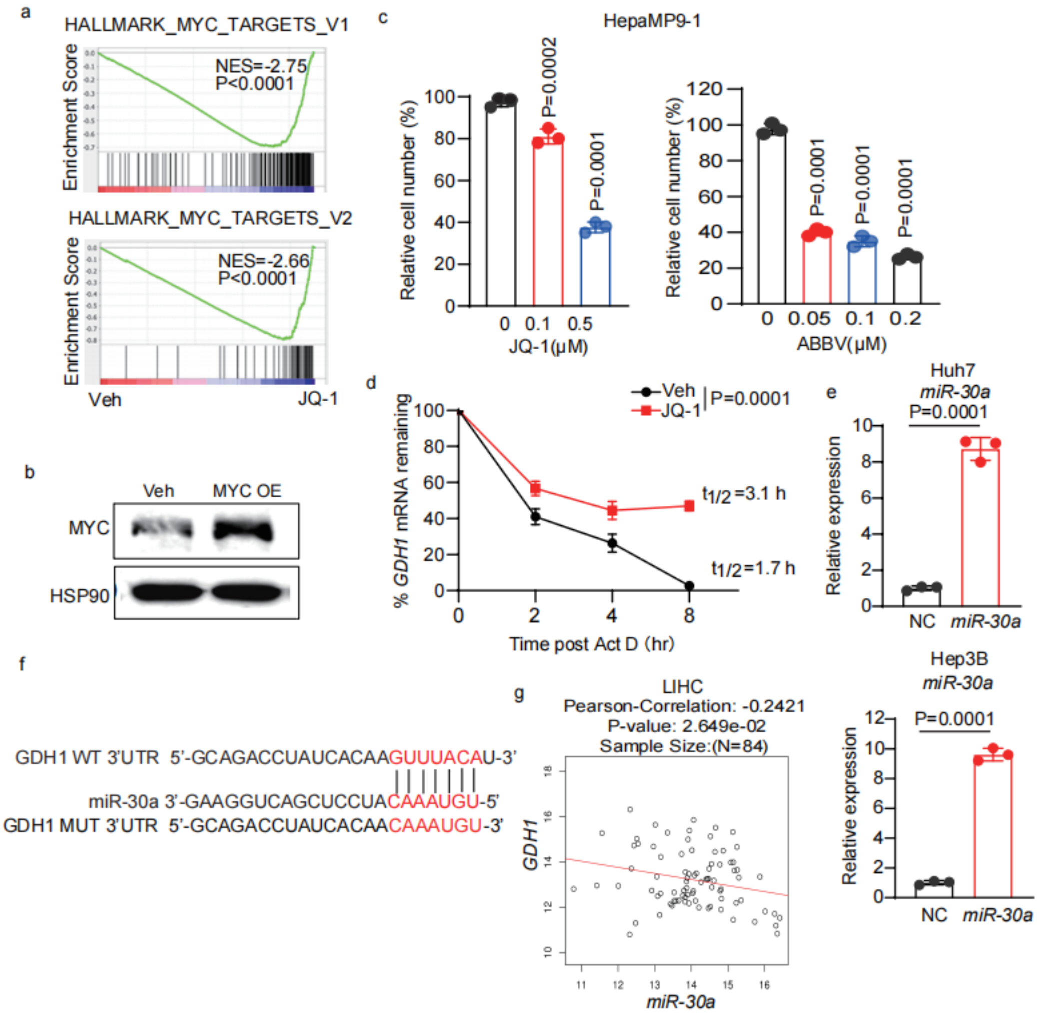


**Extended Data Fig. 5. MYC-independent GDH1 regulation by BET inhibition. a**, GSEA for MYC targets from vehicle and JQ-1 groups based on RNA-sequencing data. **b**, Western blot analysis of control and MYC-overexpressed Huh7 cells. **c**, Relative cell numbers of HepaMP9-1 cells treated with BETi for 48 hours. **d**, Q-PCR analysis of GDH1 mRNA stability in vehicle and JQ-1 treated Huh7 cells. **e**, Q-PCR analysis of *miR-30a* levels in control and miR-30a-overexpressed Huh7 and Hep3B cells. **f**, Identification of GDH1 as target by TargetScan prediction. **g**, Correlation of *miR-30a* and *GDH1* transcript levels in human HCC samples. All statistical graphs show the mean ± s.e.m. *P* values were calculated using a two-tailed Student's t-test. All experiments were performed in biological triplicate.

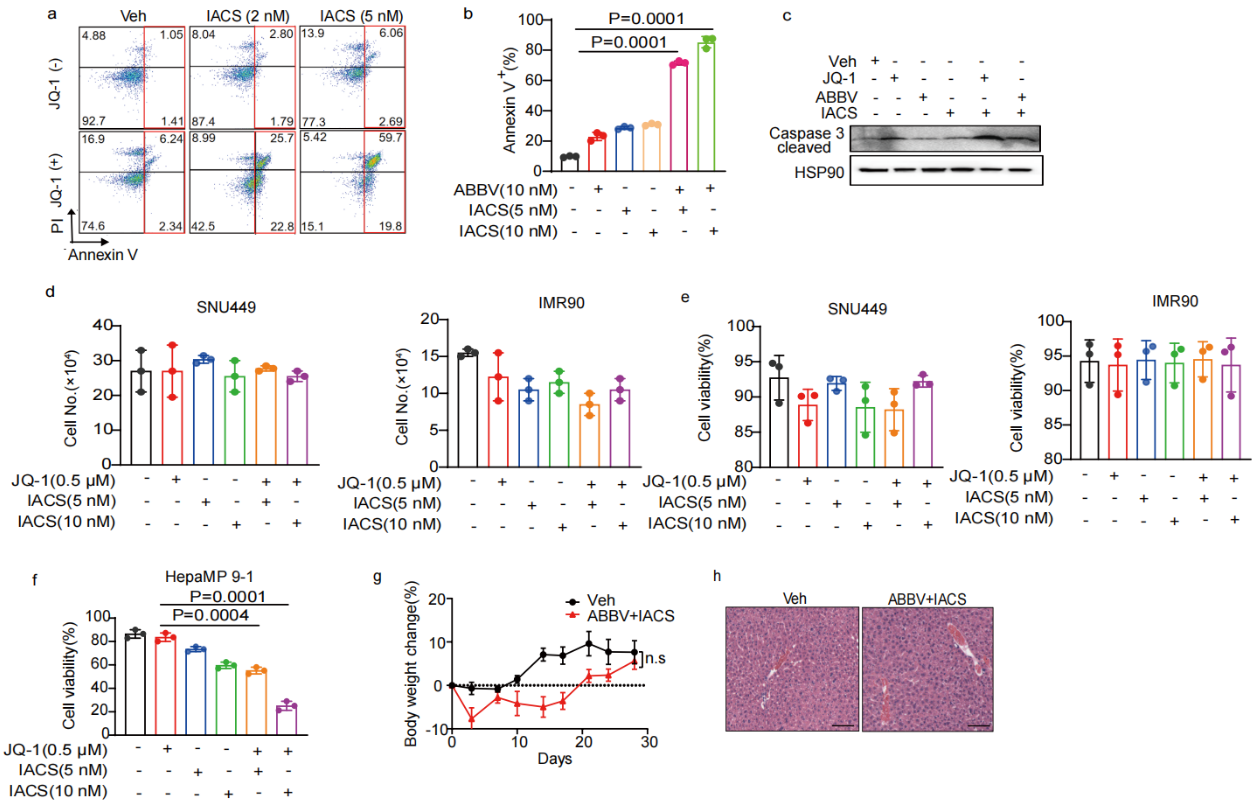


**Extended Data Fig. 6. Combined BET and OXPHOS inhibition limits liver tumor growth *in vitro* and *in vivo*. a,** Representative flow plot of PI and Annexin-V staining in the indicated JQ1 and IACS treatment groups for 6 days. **b,** Quantification of percentage of Annexin-V$^+$ Huh7 cells treated with indicated ABBV075 and IACS combination for 6 days. **c,** Western blot analysis of cleaved caspase 3 in Huh7 cells from indicated treatment groups. **d,** Quantification of the numbers of SNU449 and IMR90 cells treated with indicated drug combinations for 6 days. **e,** Quantification of the viability (% Annexin V$^-$) of SNU449 and IMR90 cells treated with indicated drug combinations for 6 days. **f**, Quantification of the viability (% Annexin V$^-$) of HepaMP9-1 cells treated with indicated drug combinations for 6 days. **g**, Body weight change over time in vehicle and combination treatment cohorts. **h**, Representative HE staining of liver sections from vehicle and combination treatment cohorts. Scale bar: 100 μm.
